# Epigenetic control and inheritance of rDNA arrays

**DOI:** 10.1101/2024.09.13.612795

**Authors:** Tamara Potapova, Paxton Kostos, Sean McKinney, Matthew Borchers, Jeff Haug, Andrea Guarracino, Steven Solar, Madelaine Gogol, Graciela Monfort Anez, Leonardo Gomes de Lima, Yan Wang, Kate Hall, Sophie Hoffman, Erik Garrison, Adam M. Phillippy, Jennifer L. Gerton

## Abstract

Ribosomal RNA (rRNA) genes exist in multiple copies arranged in tandem arrays known as ribosomal DNA (rDNA). The total number of gene copies is variable, and the mechanisms buffering this copy number variation remain unresolved. We surveyed the number, distribution, and activity of rDNA arrays at the level of individual chromosomes across multiple human and primate genomes. Each individual possessed a unique fingerprint of copy number distribution and activity of rDNA arrays. In some cases, entire rDNA arrays were transcriptionally silent. Silent rDNA arrays showed reduced association with the nucleolus and decreased interchromosomal interactions, indicating that the nucleolar organizer function of rDNA depends on transcriptional activity. Methyl-sequencing of flow-sorted chromosomes, combined with long read sequencing, showed epigenetic modification of rDNA promoter and coding region by DNA methylation. Silent arrays were in a closed chromatin state, as indicated by the accessibility profiles derived from Fiber-seq. Removing DNA methylation restored the transcriptional activity of silent arrays. Array activity status remained stable through the iPS cell re-programming. Family trio analysis demonstrated that the inactive rDNA haplotype can be traced to one of the parental genomes, suggesting that the epigenetic state of rDNA arrays may be heritable. We propose that the dosage of rRNA genes is epigenetically regulated by DNA methylation, and these methylation patterns specify nucleolar organizer function and can propagate transgenerationally.

## Introduction

Ribosome biogenesis is a fundamental housekeeping process in all cells. Ribosomal RNA (rRNA), structural non-coding RNAs, are assembled together with ribosomal proteins to generate mature ribosomes. In actively proliferating cells, most of cellular RNA production is dedicated to rRNA synthesis (Moss and Stefanovsky 2002). There are two types of ribosomal RNA: 45S rRNA and 5S rRNA. The 45S long precursor ribosomal RNA is processed through a series of cleavages and base modifications to produce 18S, 5.8S, and 28S rRNAs. These rRNAs, along with the 5S rRNA and around 80 ribosomal proteins, form the small and large ribosomal subunits (Russell and Zomerdijk 2005). 45S rRNA genes organize nucleoli, the factories for ribosome biogenesis (McStay 2016), therefore they are also referred to as nucleolar organizing regions (NORs). rRNA genes are present in multiple copies in the genomes of all eukaryotic organisms, including humans. In humans, the total copy number of 45S rRNA genes typically ranges in the hundreds and varies among individuals (Genomes Project, Auton et al. 2015, Xu, Li et al. 2017, Agrawal and Ganley 2018, Hori, Shimamoto et al. 2021). The goal of our study was to investigate the mechanism that determines the activity of rRNA gene arrays and its effect on the nucleolar organizing function.

rRNA genes are organized in tandem repeats that can be located on multiple chromosomes. In the human genome, clusters of rRNA genes are located on the short arms (p-arms) of five acrocentric chromosome pairs: 13, 14, 15, 21 and 22 (Henderson, Warburton et al. 1972). In earlier human genome assemblies, including GRCh38, the acrocentric p-arms remained as gaps because they are composed primarily of repetitive sequences, including rDNA, other segmentally duplicated genes, and various satellites. Recently, the T2T consortium used long-read sequencing techniques and innovative assembly algorithms to produce the first complete human genome assembly of the CHM13 cell line, filling gaps that had persisted for decades, including the entire acrocentric p-arms (Nurk, Koren et al. 2022). The CHM13 genome assembly showed variation in the number and sequence of 45S rRNA genes in each acrocentric array, prompting us to explore the scope of variation in rRNA gene number and activity in other human and primate genomes.

Here, we performed a comprehensive study of the number, distribution, and activity of 45S rDNA arrays across multiple human and primate genomes. We show that each individual exhibited a unique rDNA copy number, distribution of gene copies, and activity profile. Not all rDNA arrays were fully transcriptionally active, and in several cases, the entire arrays were silent. The transcriptional silencing was associated with a decreased presence of these rDNA arrays within the nucleolus, and reduced interactions with other acrocentric chromosomes, suggesting that the nucleolar organizer function is dependent on its transcriptional activity. The transcriptional silencing was associated with DNA methylation in the promoter and the coding region. Removal of this epigenetic mark restored the transcriptional activity of silent rDNA arrays. Finally, we demonstrated that an inactive rDNA array haplotype could be traced back to one of the parental genomes. This suggests that the inactive epigenetic status of rRNA genes may be heritable, being passed from parent to offspring.

## Results

### Variation in rDNA copy number and activity in human and primate genomes

The 45S rRNA genes in *Homo sapiens* are located on the short arms of acrocentric chromosomes 13, 14, 15, 21, and 22. Estimating the number of genes in each array has not been feasible using short-read sequencing, and in the earlier human genome assemblies these regions remained as gaps. The advancement of long-read sequencing technologies allowed assemblies of short rDNA arrays (Nurk, Koren et al. 2022, Rautiainen, Nurk et al. 2023, Rautiainen 2024). However, assembling and distinguishing between arrays remains challenging. To estimate the number of rRNA gene copies on each acrocentric chromosome, we used a fluorescence-based microscopy method where mitotic chromosome spreads were labeled by fluorescent in situ hybridization (FISH) with probes for rDNA and chromosome-specific markers. This allowed us to measure the fluorescent intensity of each rDNA array and assign it to a particular chromosome (Figure 1A). The fluorescence intensity of each array was quantified as a fraction of the total rDNA fluorescence signal from all arrays in the spread. This fractional value was multiplied by the total rDNA copy number estimated from Illumina sequencing to estimate the copy number of rRNA genes in each array.

**Figure 1.**
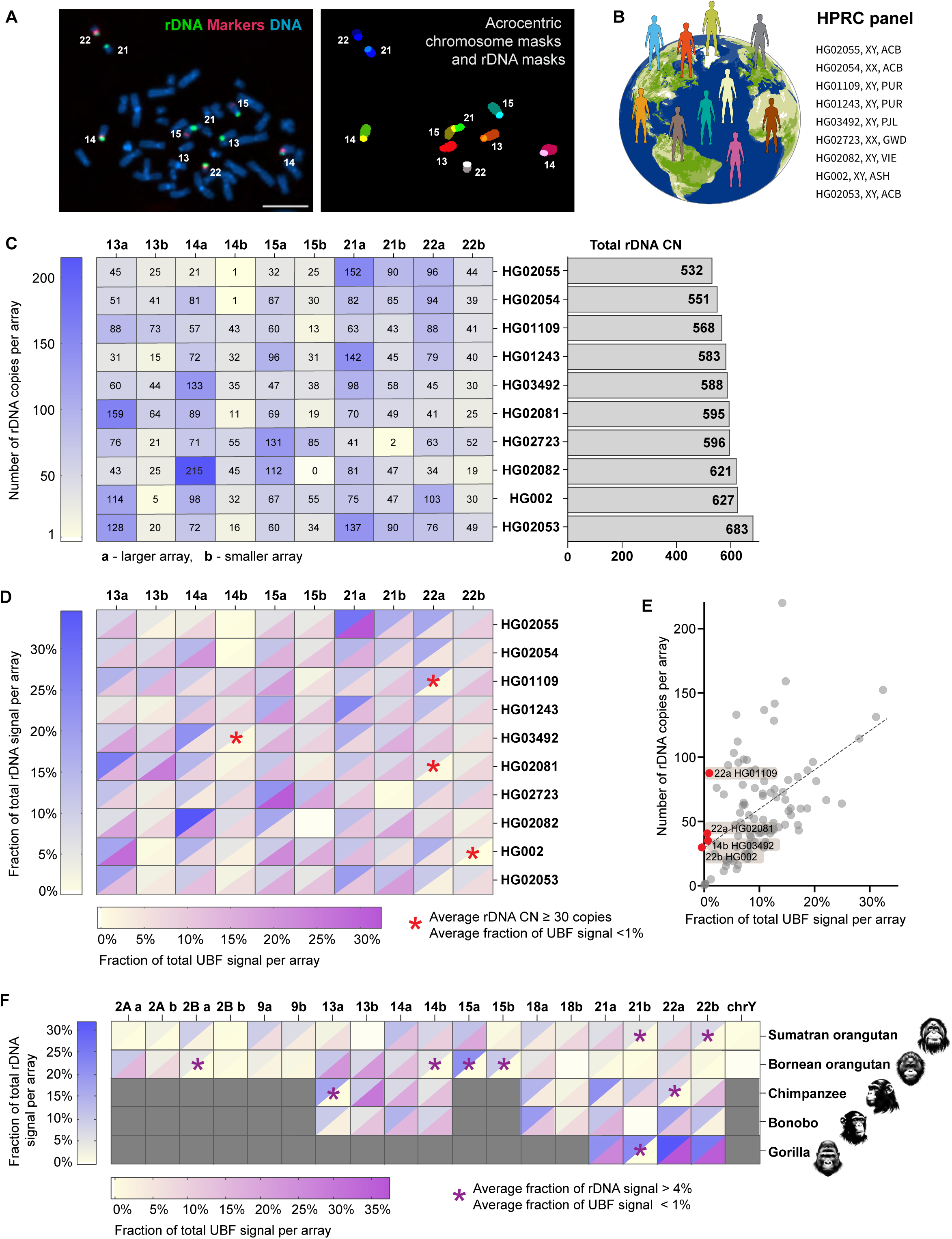
rDNA copy number and activity in human and primate genomes. **A.** Identification of specific rDNA arrays for fluorescent intensity measurements. Left panel: chromosome spread from HG002 LCL labeled by FISH with rDNA probe (green) and chromosome identification markers CenSat 14/22 and PML (red). Right panel: segmentation and identification of acrocentric chromosomes and corresponding rDNA arrays. Bar, 10µm. **B.** HPRC panel of lymphoblastoid cell lines (LCL) used in this study. **C.** Heatmap of rDNA copy numbers of each acrocentric rDNA array in the selected panel of HPRC cell lines. “a” indicates larger array and “b” smaller array. Numbers are averages from 10 or more spreads, with more detailed boxplots provided in Supplementary Figure 1. The bar plot on the right shows estimates of total rDNA CN in corresponding samples. **D.** Combined heatmap of average rDNA and UBF fluorescent intensities expressed as fractions of the total signal in a chromosome spread. rDNA was labeled by FISH, and UBF was labeled with an antibody. Both values are averages from 10 or more spreads, with detailed bar graphs shown in Supplementary Figure 2. The blue heatscale corresponds to the rDNA, and the magenta heat scale corresponds to UBF. Blank cell (15b in HG02082) indicates undetectable array. Asterisks denote arrays with 30 or more gene copies, but less than 1% of the total UBF signal. **E.** Average fractions of the total UBF signal are plotted against the average rDNA CNs for all arrays in all HPRC samples. Linear regression indicates a positive trend (R^2^=0.27). Red points denote outlier arrays with 30 or more gene copies, but less than 1% of the total UBF signal. **F.** Combined heatmap of average rDNA and UBF fluorescent intensities of rDNA arrays in non-human primates: Sumatran orangutan, Bornean orangutan, chimpanzee, bonobo, and western lowland gorilla. Chromosome identities were assigned as homo sapiens (hsa) homologues. Gray cells indicate non rDNA-bearing chromosomes in each primate species. Blank cells indicate undetectable rDNA arrays. The blue heat scale corresponds to the rDNA, and the magenta heat scale corresponds to UBF. Both parameters were expressed as percent of the total signal in a chromosome spread, showing averages from 10 or more spreads. Asterisks denote arrays with more than 4% of the total rDNA signal but less than 1% of the total UBF signal. Representative karyograms and fluorescence quantifications are shown in Supplementary Figure 3.

We developed a semi-automated pipeline using a deep learning model to perform this analysis in high throughput, and used it to investigate the rDNA copy number for a panel of ten lymphoblastoid cell lines (LCLs) derived from normal human individuals. These samples have been sequenced by the Human Pangenome Reference Consortium (HPRC) (Taylor, Eizenga et al. 2024), whose goal is to produce complete genome assemblies from a diverse cohort of human individuals (Figure 1B). The average copy number of chromosome-specific rDNA arrays showed major variation among different individuals, from very large arrays (more than 150 copies) to very small (1-2 copies) or missing arrays (Figure 1C, Sup. Figure 1A). The rDNA arrays on homologous chromosomes did not correlate in size and could contain any number of rRNA genes. Since distinguishing between maternal and paternal haplotypes was not feasible, within each homologous pair of chromosomes, the larger rDNA array was designated as “a” and the smaller as “b”. There was also no tendency of a particular acrocentric chromosome pair to contain predominantly large or predominantly small arrays. Each sample displayed a unique pattern of rDNA copy number distribution and a total rDNA copy number, suggesting that human individuals have a unique rDNA fingerprint on the p-arms of acrocentric chromosomes.

Inclusion of an rDNA activity marker allowed us to also determine the transcriptional activity of each rDNA array. rDNA has its own unique set of transcription factors, one of which is the upstream binding factor (UBF). UBF binds to the promoter and coding regions of active rRNA genes and recruits RNA Polymerase I (Pol I) (Bell, Learned et al. 1988, Mais, Wright et al. 2005). During mitosis, UBF remains associated with the rDNA, acting as a bookmark for active repeats and allowing transcription to resume in interphase (Grob, Colleran et al. 2014, Grob and McStay 2014). To measure the activity of individual rDNA arrays, chromosome spreads labeled with FISH for rDNA and chromosome identification markers were immunostained with the UBF antibody. As with rDNA copy numbers, UBF levels on chromosome-specific rDNA arrays were quantified as percentages of the total UBF signal on all arrays. Combining the averages of rDNA and UBF fractions demonstrated a unique distribution pattern of UBF in each human sample (Figure 1D, Sup. Figure 2). A general positive trend was present when all arrays were plotted together, indicating that larger arrays tend to have higher absolute levels of UBF, while smaller arrays exhibit lower UBF levels. However, a few substantial-sized arrays, containing 30 or more copies of rRNA genes, consistently had less than 1% of the total UBF signal (Figure 1E). Such arrays were identified in four out of ten HPRC samples, implying that transcriptionally silent or nearly silent rDNA arrays exist in the normal human population.

To investigate the rDNA activity status in species closely related to humans, we extended our analysis to five great ape species: Sumatran orangutan (*Pongo abelii*), Bornean orangutan (*Pongo pygmaeus*), bonobo (*Pan paniscus*), chimpanzee (*Pan troglodytes*), and western lowland gorilla (*Gorilla gorilla gorilla*). Chromosome spreads were obtained from primary male fibroblast and lymphoblastoid cell lines from these species, one from each individual animal. To identify primate rDNA-bearing chromosomes we used human whole-chromosome paints that allowed us to distinguish *Homo sapiens* (hsa) analogues. Genomes of apes retain the ancestral pre-fusion state of human chromosome 2: hsa 2A and 2B (Yunis and Prakash 1982). Orangutans have the highest number of acrocentric chromosomes containing rDNA arrays, that is nine pairs: 2A, 2B, 9, 13, 14, 15, 18, 21, and 22 (Chiatante, Giannuzzi et al. 2017), as well as chromosome Y (Makova, Pickett et al. 2024). These arrays varied in size, and multiple arrays were UBF-negative in both species (Figure 1F, Sup. Figure 3 A, B). One copy of chromosome 13 in Sumatran orangutan and one copy of chromosome 18 in Bornean orangutan were missing rDNA arrays. UBF was indiscernible on chromosome Y in Sumatran orangutan, which may be due to its small rDNA array size (3 copies), and the single chromosome Y rRNA gene in Bornean orangutan was undetected by FISH in our experiments. Chimpanzees and bonobos have rRNA genes on five pairs of acrocentric chromosomes (13, 14, 18, 21, and 22). In our chimpanzee specimen, two large arrays on chromosomes 13 and 22 were UBF-negative (Figure 1F, Sup. Figure 3 C). In the bonobo specimen, all rDNA arrays were UBF-positive, but one array on chromosome 21 was missing (Figure 1F, Sup. Figure 3 D). Gorillas have only two pairs of rDNA-bearing chromosomes (21 and 22). Still, one of the arrays on chromosome 21, which comprised approximately 22% of the total rRNA gene pool, was negative for UBF in our specimen (Figure 1F, Sup. Figure 3 E). These results demonstrate that the copy number and activity variation of rDNA arrays are not unique to humans and may be a common feature across primate genomes.

### The activity status of rDNA arrays determines their nucleolar organizing function

We further explored rDNA activity in the CHM13 cell line, which was the source of the first complete telomere-to-telomere (T2T) human genome assembly (Nurk, Koren et al. 2022). This cell line, derived from an abnormal gestational tissue called hydatidiform mole, possesses a uniparental homozygous genome where both sets of homologous chromosomes are paternal and an epigenetic profile similar to a trophoblast (Carey, Nash et al. 2015, Huddleston, Chaisson et al. 2017). The diploid and mostly homozygous genome of the CHM13 cell line allowed us to explore rDNA activity status and spatial organization without the confounding factors of haplotype variation. rDNA arrays on homologous pairs were combined and averaged since they were homozygous, with the exception of chromosome 15, where the arrays were different sizes due to a heterozygous deletion in the cell line (Nurk, Koren et al. 2022). Results from UBF immuno-FISH experiments showed that in CHM13 cells, the rDNA arrays on both copies of chromosome 22 were UBF-negative, indicating that chromosome 22 rDNA arrays were transcriptionally silenced (Figure 2A). This observation was validated by performing immuno-FISH with antibodies against Treacle, another component of the rDNA transcriptional machinery that can serve as an indicator of rDNA transcription (Valdez, Henning et al. 2004). The UBF-negative rDNA arrays on chromosome 22 were also negative for Treacle (Figure 2B), confirming that the rDNA arrays on this chromosome pair are transcriptionally inactive in the CHM13 cell line.

**Figure 2.**
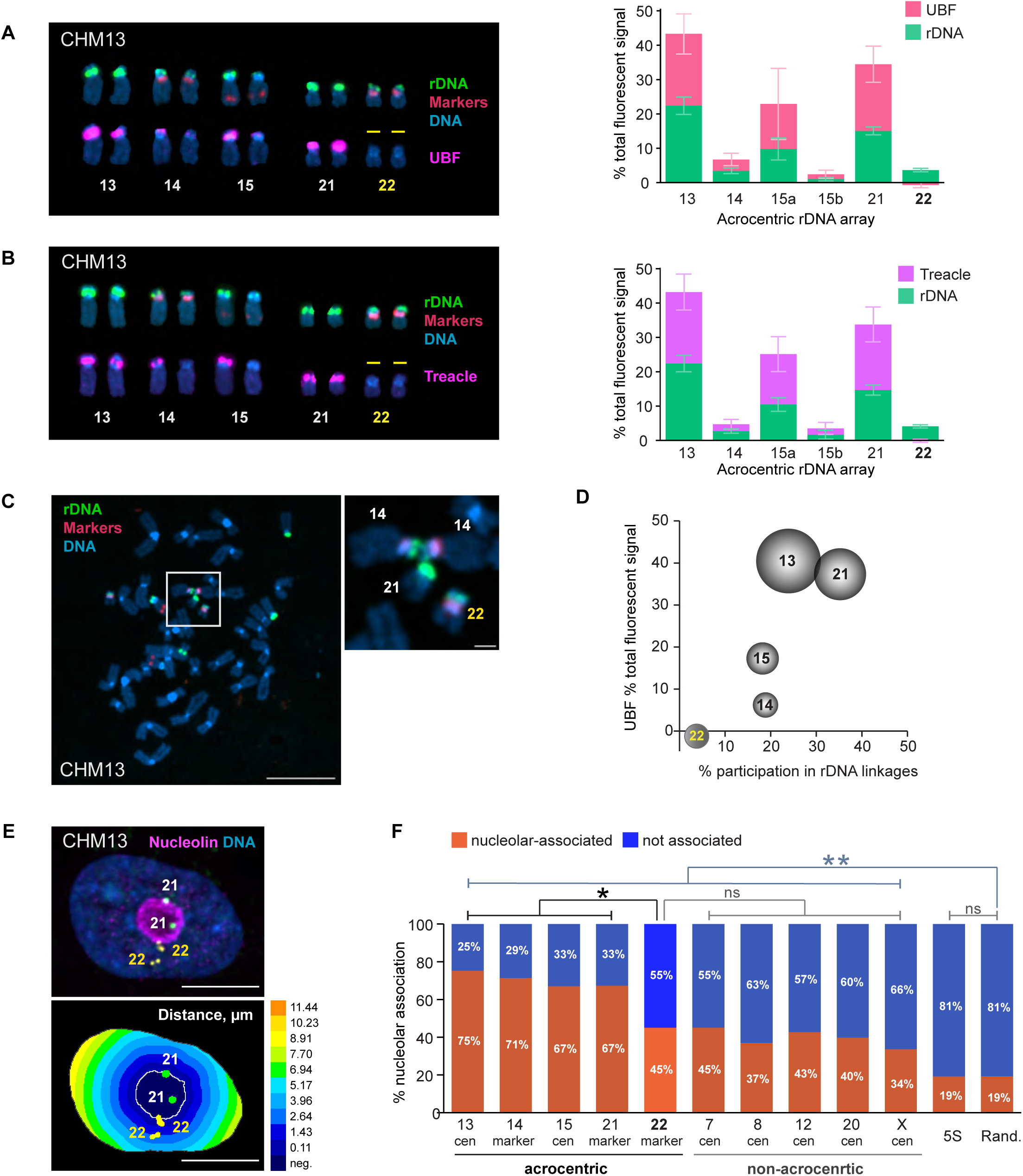
Effects of rDNA activity status on 3D organization in CHM13 cell line. **A.** Right panel: acrocentric karyogram from a representative CHM13 chromosome spread labeled by immuno-FISH with rDNA probe and UBF antibody. Top row of chromosomes shows FISH labeling with rDNA probe (green) and chromosome identification markers CenSat 14/22 and PML (red). Bottom row shows corresponding chromosomes with UBF antibody labeling (magenta). DNA was counter-stained with DAPI. Note that both rDNA arrays on chromosome 22 are UBF-negative. Left panel: Quantification of rDNA FISH and UBF antibody labeling on acrocentric arrays. rDNA FISH and UBF antibody signals were measured as fractions of the total fluorescent intensity in the chromosome spread. Since CHM13 cell line is homozygous diploid, both homologous acrocentric arrays were averaged except for chromosome 15. The pink and green sections of the bars represent averages of UBF and rDNA, respectively. Error bars denote standard deviation. **B.** Treacle antibody was used for rDNA activity estimation in CHM13 by immuno-FISH and as in A. Note that both rDNA arrays on chromosome 22 are also Treacle-negative. **C.** A representative chromosome spread from CHM13 cell line showing an example of the rDNA linkage between heterologous acrocentric chromosomes. rDNA (green) and chromosome identification markers (red) were labeled by FISH, DNA was counter-stained with DAPI. Bar, 10µm. Magnified insert shows the rDNA linkage between two copies of chromosome 14 and a copy of chromosome 21. Bar, 1µm. **D.** Relationship between the frequency of rDNA linkages and the activity of rDNA arrays measured by UBF as fractions of the total fluorescent signal. The percent participation in rDNA linkages was determined as the fraction of total linkage occurrences for a particular chromosome. The dimensions of the spheres reflect sizes of rDNA arrays. Both homologous acrocentric arrays were averaged. Large and highly active rDNA arrays (chromosomes 13 and 21) formed linkages frequently, while the inactive arrays on chromosome 22 very rarely participated in linkages. **E.** Top panel: immuno-FISH image of a representative CHM13 nucleus labeled with rDNA-adjacent markers for chromosome 21 (green), chromosome 22 (yellow), and nucleoar marker nucleolin (magenta). Nuclei were counter-stained with DAPI (blue). Bar, 10µm. Bottom panel: Euclidean Distance Transform (EDT) map of the same nucleus with nucleolus and near-centromeric markers segmented. The intensity scale indicates the distance to nucleolar boundary, µm. **F.** Nucleolar association of rDNA-adjacent acrocentric chromosome-specific markers, non-acrocentric centromeric markers, and 5S rDNA loci in CHM13 cells. The orange and blue sections of the bars represent fractions of nucleolar-associated and not associated loci, respectively. Validation of rDNA-adjacent chromosome-specific markers is shown in Supplementary Figure 4A. The nucleolar association of chromosome 22 marker is significantly reduced compared to the active acrocentric chromosome markers and is not significantly different from non-acrocentric centromeric markers (Kolmogorov-Smirnov test). Note that nucleolar association of both acrocentric and non-acrocentric markers was significantly higher than that of 5S rDNA locus or random points in the nucleus. Distributions of distances from the nucleolar boundary for all markers are shown in Supplementary Figure 4B.

rRNA gene arrays organize the nucleoli during interphase (McStay 2016, Potapova and Gerton 2019). Within the shared nucleolar compartment arrays of rRNA genes can engage in physical interchromosomal associations. We have previously shown that rDNA transcription can lead to the formation of topological intertwines, or linkages, between transcriptionally active arrays from heterologous chromosomes, that are mediated and resolved by Topoisomerase II (Potapova, Unruh et al. 2019). Examining rDNA linkages in CHM13 cells supported our hypothesis about the role of rDNA transcription in the formation of these interchromosomal interactions, first described by Ferguson-Smith in 1961 (Ferguson-Smith and Handmaker 1961). We quantified the frequency of rDNA linkages as the percent of total linkage occurrences for a particular chromosome. Larger rDNA arrays with high activity, such as those on chromosomes 13 and 21, formed linkages more frequently (24% and 35%, respectively). However, chromosome 22 arrays rarely (4%) participated in rDNA linkages (Figure 2 C-D). Therefore, transcriptional silencing of the rDNA reduces its intertwining with other, active arrays, and reduces interchromosomal interactions with other NOR-bearing chromosomes.

Next, we evaluated the nucleolar organizing capability of the transcriptionally silent chromosome 22 rDNA array. The nucleolar compartment assembles in interphase around transcriptionally active rDNA (Hernandez-Verdun 2011). Its assembly is mediated by specific interactions of nucleolar proteins with rDNA and rRNA, as well as liquid-liquid phase-separation forces (Hori, Engel et al. 2023). Additionally, genomic regions other than rDNA have also been shown to play a role in the assembly of functional nucleolar compartments. For instance, the rDNA distal junction (DJ) sequence located distally to the rDNA arrays in humans, was proposed to have an anchoring function (van Sluis, Gailin et al. 2019, Liskovykh, Petrov et al. 2023). Furthermore, centromeres of NOR-bearing and other chromosomes were shown to be associated with the nucleolus in various model systems (Haaf and Schmid 1989, Ochs and Press 1992, Carvalho, Pereira et al. 2001) and potentially play a role in nucleolar organization (Rodrigues, MacQuarrie et al. 2023). The T2T assembly of the CHM13 genome did not indicate any alterations of the DJ region or other parts of the acrocentric p-arms of chromosome 22. Thus, while the transcriptional activity of chromosome 22 rDNA arrays was silenced, their genomic context was normal. In the interphase nucleus, it was not possible to assign the rDNA signal to specific chromosome arrays because the rRNA genes are indistinguishable by FISH and intertwined within the nucleoli. To overcome this, we used centromeric or near-centromeric markers specific to each chromosome located in close proximity to the rDNA (Sup. Figure 4A). We then measured the distance from these markers to the nearest nucleolus using Euclidean distance transform (EDT) maps (Figure 2E). Setting the threshold for nucleolar association at 0.5 µm from the nearest nucleolar boundary, we measured the fractions of rDNA-adjacent markers associated and not associated with the nucleoli. For acrocentric chromosomes with active rDNA, the nucleolar association rate ranged from 67% to 75%. In contrast, chromosome 22 exhibited a significantly lower association rate of 45%, comparable to centromeres of non-acrocentric chromosomes that do not contain any rDNA or DJ sequences. These rates of nucleolar association were all significantly higher than those of the 5S locus located at the end of the long arm of chromosome 1 or randomly positioned points within the nucleus (Figure 2F, Sup. Figure 4B). These results suggest that while inactive rDNA arrays may not function as nucleolar organizing regions, they can still associate with the nucleoli at a rate comparable to the centromeric regions of non-rDNA chromosomes. Notably, centromeres of non-acrocentric chromosomes showed a higher-than-random association with the nucleolus, indicating a propensity to associate with perinucleolar heterochromatin irrespective of rDNA.

### rRNA genes are epigenetically silenced by DNA methylation

To investigate transcriptional silencing of the rDNA array on chromosome 22, we examined its methylation status in CHM13 cells. Each human rDNA repeat unit is composed of the promoter, the coding region for 18S, 5.8S and 28S rRNA molecules separated by internal transcribed spacers and flanked by external transcribed spacers, and followed by the non-transcribed intergenic spacer (IGS) (Kim, Dilthey et al. 2018). DNA methylation is an important regulator of gene expression in mammals (Schubeler 2015). Previous analysis of long-read sequencing data from multiple human genomes demonstrated that the rDNA promoter and the coding region generally existed in two distinct epigenetic states—unmethylated and methylated, while IGS was methylated in both states (Hori, Shimamoto et al. 2021). A remaining challenge is assigning rDNA reads to specific chromosomes. We approached chromosome assignment by isolating individual acrocentric chromosomes from mitotic CHM13 cells by fluorescence-activated cell sorting (FACS) (Figure 3A). DNA from sorted chromosome populations was analyzed by an enzymatic methyl-sequencing method, or methyl seq, an alternative to conventional bisulfite sequencing. This technique converts unmethylated cytosines to uracils enzymatically, minimizing the noise typically caused by the bisulfite conversion reaction (Vaisvila, Ponnaluri et al. 2021). FACS-sorting of acrocentric chromosomes followed by DNA sequencing verified that most of the reads were derived from specific acrocentric chromosome populations (Figure 3B).

**Figure 3.**
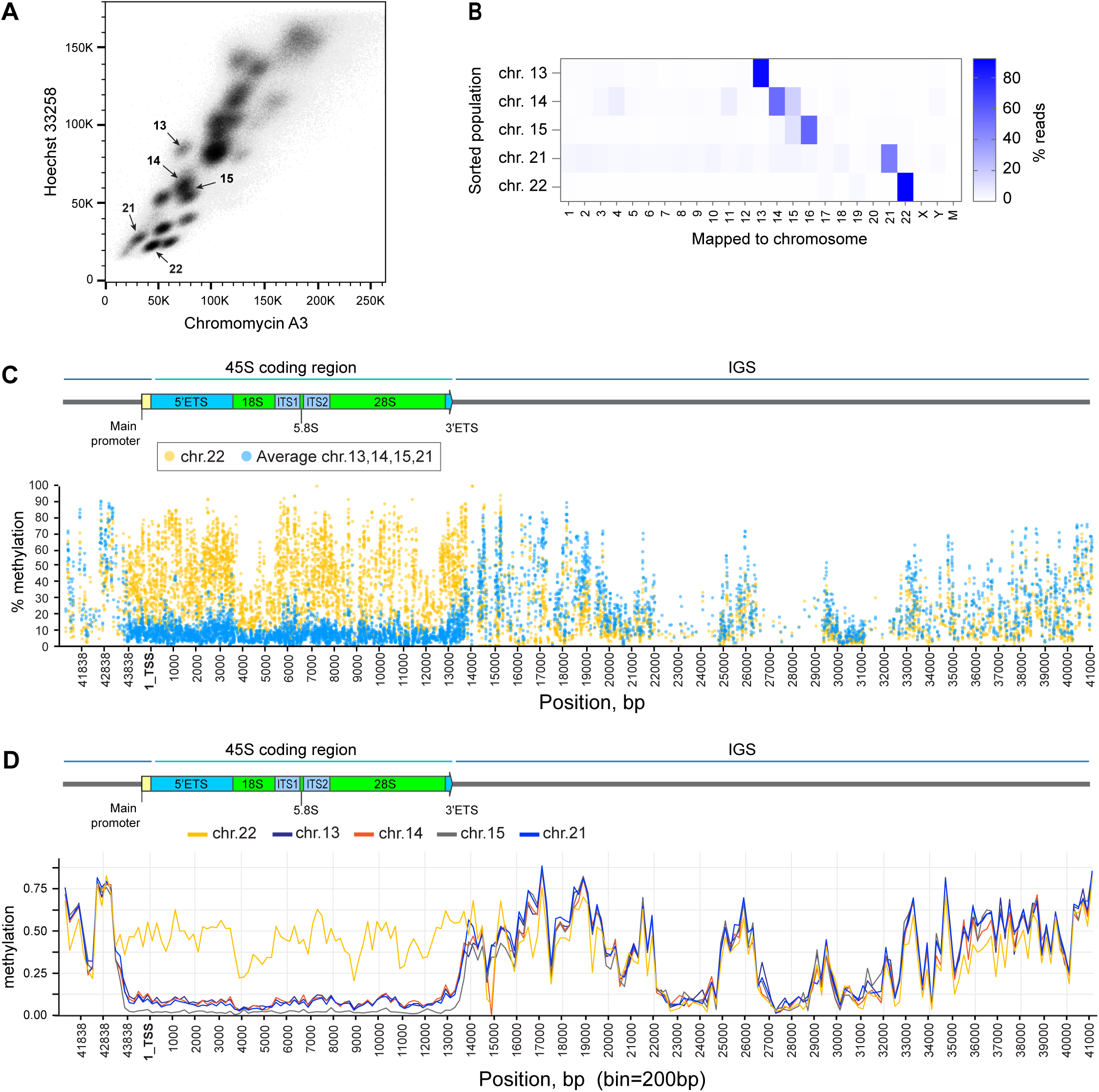
Epigenetic silencing of rDNA by methylation in CHM13 cell line. **A.** Acrocentric chromosome sorting by flow cytometry. Chromosomes were isolated from CHM13 cell line as detailed in the Materials and Methods and labeled with Chromomycin A3 and Hoechst 33258. Arrows indicate sorted populations. **B.** Illumina short read mapping from flow-sorted chromosomes. The purity of sorted populations was confirmed by the highest fraction of reads mapped to expected chromosomes, indicated by the color scale. **C.** Percent methylation of each cytosine base in reads mapped to the rDNA reference sequence across the rRNA gene determined by short-read methyl-sequencing analysis. The promoter and the coding region of transcriptionally inactive chromosome 22 rDNA (yellow circles) are highly methylated compared to averages of all other acrocentric chromosomes with transcriptionally active rDNA (blue circles). Individual plots for each chromosome are shown in Supplementary Figure 5 A-D. **D.** Methylation calls from ONT long-read sequencing of rDNA reads mapped to specific acrocentric chromosomes. Reads mapped to chromosome 22 (yellow line) are highly methylated in the promoter and coding region compared to reads mapped to other acrocentric chromosomes. Bin size 200bp.

The methyl-sequencing analysis showed a higher percentage of cytosine methylation in the promoter and coding regions of rDNA reads from silent chromosome 22 arrays, compared to the average methylation levels of reads from all other, active arrays at the same positions at a single-base resolution (Figure 3C). The methylation patterns of rDNA reads from active arrays were very similar, displaying a uniform drop in cytosine methylation starting around 1kb upstream of the promoter region and continuing throughout the promoter and the coding region (Sup. Figure 5). The intergenic spacer was highly methylated in all arrays, and the methylation patterns in this region were comparable across different sorted chromosomes. These results were further confirmed by analyzing methylation calls from Oxford Nanopore (ONT) long-read sequencing data used for the CHM13 T2T assembly. In the case of CHM13, the individual rDNA arrays are distinct enough that individual ONT reads could be assigned to a particular acrocentric chromosome based on their sequence and alignment to the reference assembly (Nurk, Koren et al. 2022). The methylation landscape of these chromosome-assigned ONT reads across the rDNA repeat unit also showed higher methylation levels in the promoter and coding region of chromosome 22 reads compared to all other chromosomes (Figure 3D). These findings revealed that the transcriptionally inactive rDNA on chromosome 22 in CHM13 cells is highly methylated across the entire unit.

To confirm that methylation of the rDNA promoter and coding region was responsible for silencing RNA Pol I transcription, we performed a gain-of-function experiment using an inhibitor of DNA methyltransferase DNMT1. DNMT1 maintains cytosine methylation patterns throughout the genome by conversion of cytosine residues to 5-methylcytosines during DNA replication (Hermann, Goyal et al. 2004). We utilized a selective DNMT1 inhibitor GSK-3484862 that was shown to globally reduce DNA methylation after prolonged treatments, with low cytotoxicity (Azevedo Portilho, Saini et al. 2021). CHM13 cells were cultured in the presence of 5µM GSK-3484862 for one month. Subsequently, the inhibitor was washed out, cells were allowed to recover in drug-free medium and analyzed by whole genome methyl-sequencing and UBF immuno-FISH. Methyl-sequencing demonstrated that in drug-treated cells methylation of rDNA reads was strongly reduced across all regions, including the promoter, the coding region, and the IGS (Figure 4A). Analysis of all DNA reads mapped to the entire acrocentric chromosomes confirmed that the drug caused cytosine de-methylation genome-wide (Sup. Figure 6A). Immuno-FISH analysis showed that in the GSK-3484862 treated population of cells, chromosome 22 rDNA arrays became positive for UBF in 75% of chromosome spreads following this treatment (Figure 4B). In some instances, only one array was UBF positive (20% of cases), while in others both arrays were positive (55% of cases) with varying degrees of UBF intensities (Figure 4C, Sup. Figure 6B). The transcriptionally silent rDNA array on chromosome 22 was reactivated to some degree in the majority of cells treated with DNMT1 inhibitor, implying that rDNA methylation is required to maintain its silenced state.

**Figure 4.**
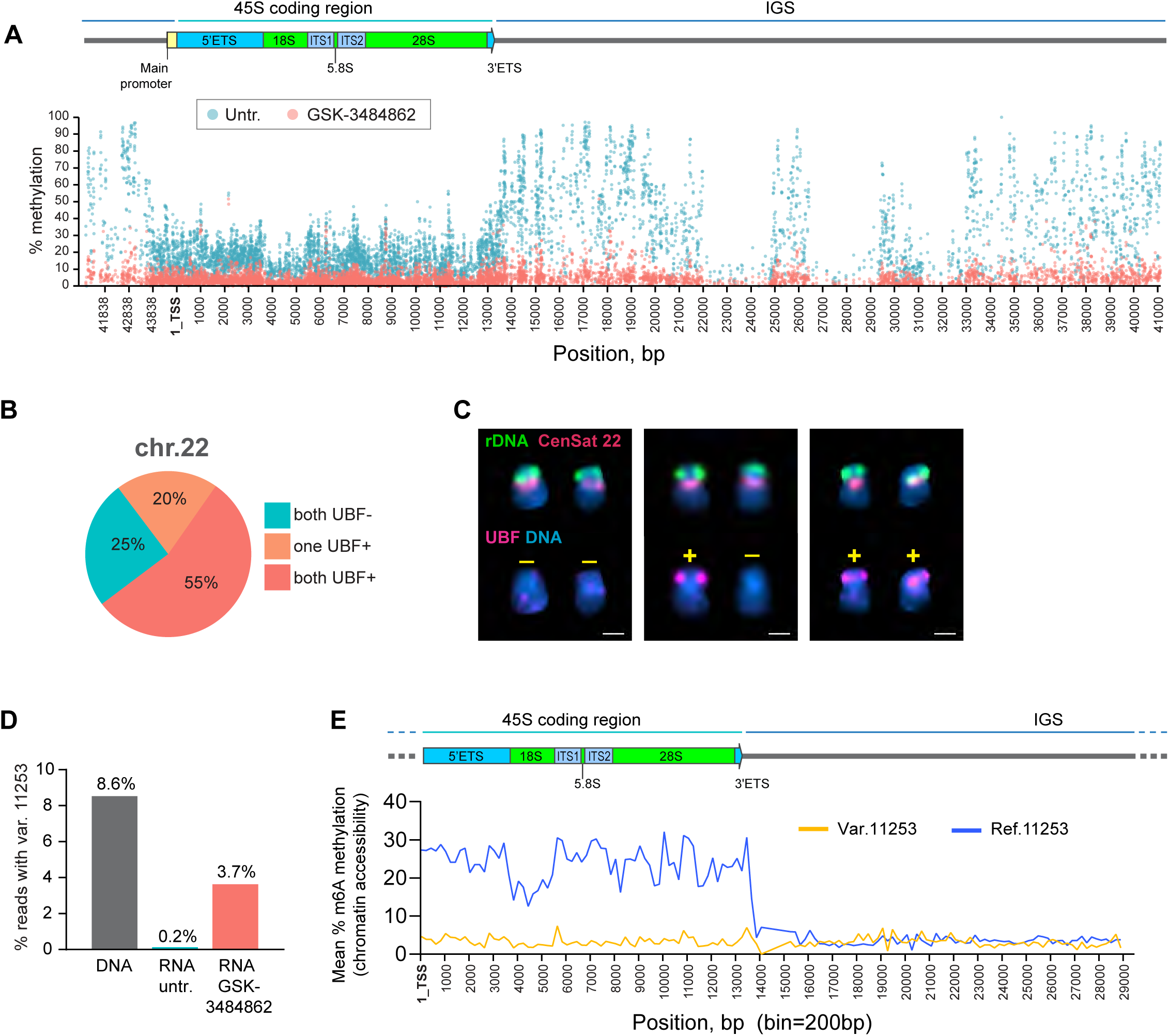
DNMT1 inhibitor restores the activity of silent rDNA arrays in CHM13 cell line. **A.** Methyl-sequencing analysis of CHM13 cells grown in the presence of the DNMT1 inhibitor GSK-3484862 for four weeks, with an untreated culture maintained in parallel. The percent methylation of each cytosine base in reads mapped to the rDNA reference sequence across the entire rRNA gene is shown. The DNMT1 inhibitor reduces DNA methylation levels throughout the gene, including the promoter, coding region, and intergenic spacer. **B.** UBF immuno-FISH results from forty chromosome spreads of CHM13 cells treated with the DNMT1 inhibitor. The fractions of spreads where one, both, or neither rDNA arrays on chromosome 22 re-gained UBF signal are shown. **C.** Examples of chromosome 22 pairs from UBF immuno-FISH experiment show panel in B. The top rows of chromosomes show FISH labeling with rDNA probe (green) and CenSat 22 (red). The bottom rows show corresponding chromosomes with UBF antibody labeling (magenta). DNA was counter-stained with DAPI. UBF status is indicated by +/-. Complete karyograms are provided in Supplementary Figure 6B. **D.** Fractions of rRNA reads containing a variant at position 44997. RNA from untreated and GSK-3484862-treated cells was used to create non-ribodepleted libraries. Illumina short sequencing reads were analyzed using the mpileup tool to detect the presence of the variant at position 44997. DNA from the CHM13 cell line was analyzed in parallel. The DNMT1 inhibitor treatment caused re-expression of the silent variant. **E.** Fiber-seq mean percent methylation of adenine bases in rDNA-containing reads. Reads containing chromosome 22-specific variant at position 11253 (yellow line) are less methylated in the coding region compared to reads containing the reference base (blue line), indicating less accessible chromatin. Bin size 200 bp.

Human rRNA genes exhibit sequence variation, predominantly in the IGS, although the coding regions, particularly the external and internal transcribed spacers and the 28S region, also contain sequence variants (Kim, Dilthey et al. 2018, Fan, Eklund et al. 2022). The variants in the coding region may be expressed in the rRNA pool, or they may be silent (Rothschild, Susanto et al. 2024). The chromosome 22-specific SNP variant at the position 11253 (A->G) in the 28S region was detected in the DNA but not in the rRNA of untreated cells (Figure 4D). A separate chromosome sorting experiment followed by the rDNA variant calling demonstrated that the majority of the 28S rDNA reads containing this SNP were present on chromosome 22 (Sup. Figure 6C). In drug-treated cells, this variant became detectable in the 28S rRNA (Figure 4D), confirming that removing rDNA methylation promotes the reactivation of the silent chromosome 22 rDNA array and expression of this variant. Next, we examined chromatin accessibility of the rDNA arrays in CHM13 by the Fiber-seq technique (Dubocanin D 2024, Jha, Bohaczuk et al. 2024). Fiber-seq is a long read sequencing method that maps chromatin accessibility by the deposition of m6A along open DNA (Stergachis, Debo et al. 2020). PacBio HiFi Fiber-seq reads containing the 28S SNP variant at the position 11253 had significantly lower m6A methylation along the gene body compared to reads that did not contain this variant (Figure 4E), indicating a closed chromatin environment characteristic of silent genes.

Our interpretation of these results is that rDNA methylation constitutes one of the mechanisms controlling the dosage of active rRNA genes by keeping some of the repeats silent, or, in the case of CHM13, by silencing entire arrays. This finding aligns with the positive correlation reported by Hori et al. (Hori, Shimamoto et al. 2021) between the number of methylated rRNA genes and the total number of rRNA genes. Therefore, methylation of the 45S genes and closed chromatin conformation may be a defining characteristic of the commonly observed UBF-negative arrays in humans and non-human primates.

### rDNA activity status may be heritable

To extend our observations in the CHM13 cell line to diploid human individuals, we investigated rDNA activity in the HG002 genome. The HG002 genome, a heterozygous diploid assembly from a human male, is a high-quality reference genome, although rDNA regions still contain gaps at this time (Jarvis, Formenti et al. 2022). Our UBF immuno-FISH results from the HG002 lymphoblastoid cell line (LCL), obtained during the initial HPRC panel analysis, identified an inactive array on one of the copies of chromosome 22 (Figure 1 D-E, Sup. Figure 2). For the HG002 genome, induced pluripotent stem cells (iPSCs) and LCLs from both maternal and paternal individuals are available, allowing us to investigate reprogramming and heritability of rDNA activity status.

First, we tested whether the silent rDNA array was re-activated by stem cell reprogramming. Epigenetic reprogramming is a necessary process in generation of induced pluripotent stem cells. DNA of undifferentiated iPS cells is generally hypomethylated (De Carvalho, You et al. 2010, Nishino, Toyoda et al. 2011), although some epigenetic memory can persist (Kim, Doi et al. 2010). We obtained two iPS cell lines derived from the HG002 B-lymphocytes, designated iPS1 and iPS2, and cultured them in feeder-free conditions. These isogenic cell lines differed in the delivery mode of the reprogramming factors: iPS1 was re-programmed using episomal delivery, and the iPS2 using the Sendai virus. Surprisingly, the UBF pattern in both iPS cell lines was similar to that of the LCL across most acrocentric arrays (Figure 5A). The sizes of rDNA arrays also remained the same on most chromosomes. In HG002, the chromosome 22 pair carries two distinct rDNA array haplotypes: a larger, UBF-positive array (“a”) and a smaller, UBF-negative array (“b”). The smaller rDNA array was persistently UBF-negative in LCLs and in both iPS cell lines (Figure 5B, Sup. Figure 7A), indicating it was not re-activated by reprogramming.

**Figure 5.**
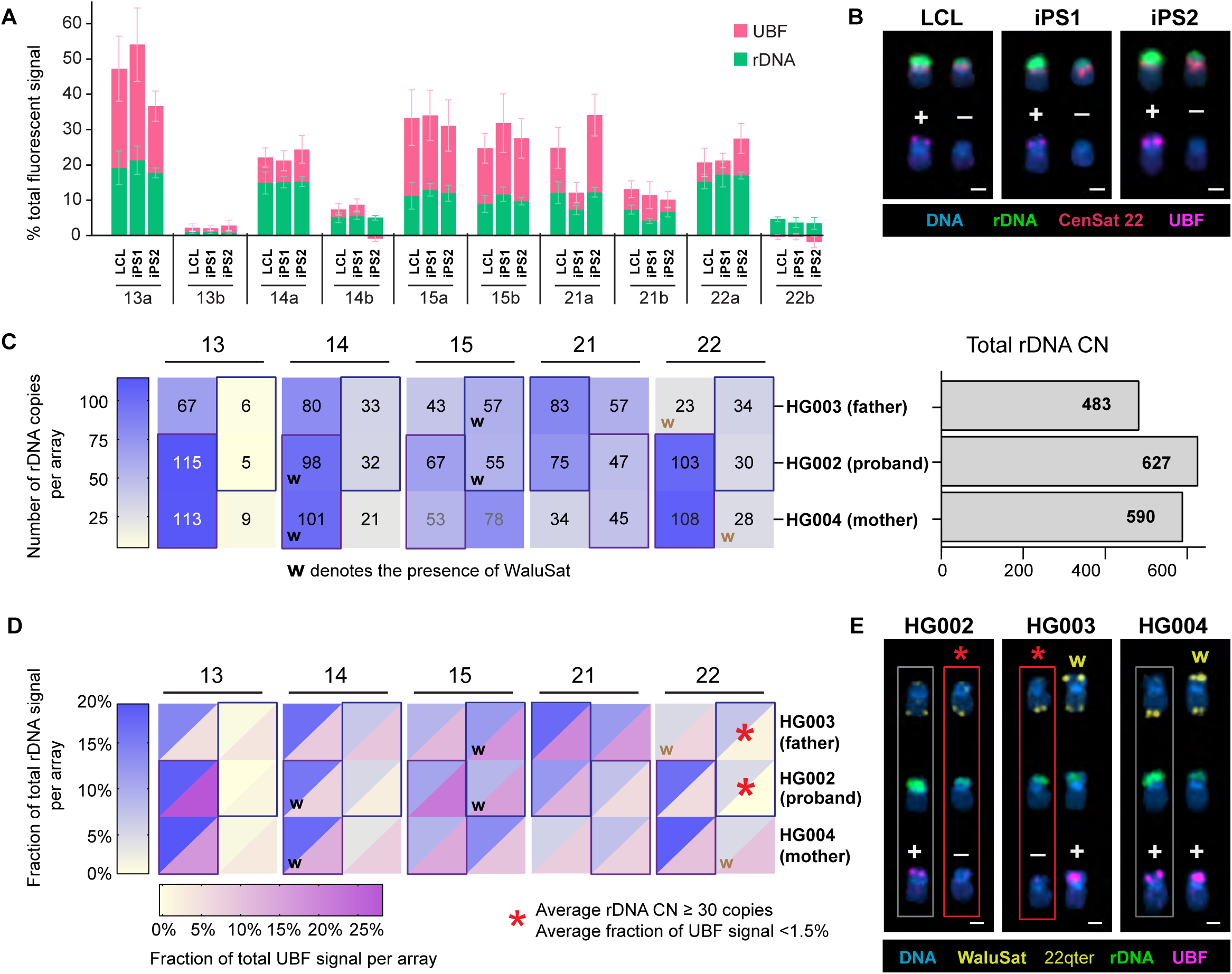
rDNA copy number and activity status in HG002 iPS cells and family trio. **A.** Quantification of rDNA FISH and UBF antibody labeling on acrocentric arrays in HG002 LCL and two iPS cell lines. The rDNA FISH and UBF antibody signals were measured as fractions of the total fluorescent intensity in the chromosome spread. Note the smaller rDNA array on chromosome 22 that is UBF-negative in all three cell lines. The green and pink sections of the bars represent the averages of rDNA and UBF signals, respectively. Error bars denote standard deviations. **B.** Examples of chromosome 22 pairs from HG002 cell lines measured in panel A. The top rows display FISH labeling with the rDNA probe (green) and CenSat 22 (red). The bottom rows show corresponding chromosomes with UBF antibody labeling (magenta). DNA was counterstained with DAPI. UBF status is indicated by +/-. Extended karyograms are provided in Supplementary Figure 7A. **C.** Heatmap of rDNA copy numbers for each acrocentric rDNA array in the HG002 family trio. The proband’s arrays are sorted by size (large and small), while parental arrays are arranged to match the proband. Boxes indicate approximations for inherited arrays, and “w” denotes the presence of WaluSat. Numbers represent averages from 10 or more spreads, with detailed boxplots in Supplementary Figure 1 for HG002, and Supplementary Figure 6B for HG003 and HG004. The bar plot on the right estimates the total rDNA copy number in corresponding samples. **D.** Combined heatmap of average rDNA and UBF fluorescent intensities, expressed as fractions of the total signal in a chromosome spread. Both values are averages from 10 or more spreads, with detailed bar graphs provided in Supplementary Figure 2 for HG002, and Supplementary Figure 7C for HG003 and HG004. The blue heat scale corresponds to rDNA, and the magenta heat scale corresponds to UBF. The asterisk denotes the low activity rDNA array present on the WaluSat-negative copy of chromosome 22 in HG002 (proband) and HG003 (father). **E.** Examples of chromosome 22 pairs from HG002, HG003, and HG004 LCL cell lines shown in panel D. The top rows display chromosomes labeled by FISH with WaluSat probe (highlighted by arrows) and a chromosome 22 q-terminal region probe (arrow) serving as an identification marker. The middle rows show corresponding chromosomes labeled with the rDNA probe (green). The bottom rows display the corresponding chromosomes labeled with UBF antibody (magenta). DNA was counterstained with DAPI. The UBF-negative rDNA array on the WaluSat-negative copy of chromosome 22 in HG002 and the corresponding paternal copy of chromosome 22 in HG003 are highlighted by red boxes and asterisks. The maternal copy of chromosome 22 is highlighted by gray

Next, we explored the epigenetic status of the inactive chromosome 22 rDNA array in the parental genomes. For the HG002 individual, family trio LCL cell lines are available, derived from HG003 (father) and HG004 (mother). These cell lines were used for the cytogenetic paternity analysis with quantification of rDNA array size as a marker of paternity for acrocentric chromosomes. As a confirmatory marker, we used an acrocentric p-arm distal satellite termed WaluSat. WaluSat is a human satellite present exclusively on the short arms of some, but not all, acrocentric chromosomes distal to the rDNA (Hoyt, Storer et al. 2022, Nurk, Koren et al. 2022). Its presence on specific acrocentric chromosomes varies among individuals. The results show that the acrocentric haplotypes in the HG002 individual can be traced back to both the father and the mother, based on the sizes of the rDNA arrays and the presence of the WaluSat marker (Figure 5C, Sup. Figure 1, 7B). In both paternal and maternal genomes, WaluSat was present at detectable levels on one of the copies of chr.22, neither of which was inherited by the proband. UBF immuno-FISH analysis showed that the activity status also generally propagated with parental haplotypes (Figure 5D, Sup. Figure 2, 7C). The large UBF-positive rDNA array on the maternal WaluSat-negative copy of chromosome 22 remained UBF-positive in the proband. The paternal rDNA array on the WaluSat-negative copy of chr.22 inherited by the proband had extremely low UBF signal (Figure 5E, Sup. Figure 7D). Assuming no recombination, the low activity status of rDNA array on chr.22 propagated from father to child. This suggests that epigenetic rDNA silencing may propagate through generations, maintaining rRNA genes or entire arrays in a transcriptionally inactive state.

## Discussion

rDNA has been extensively studied for a long time, and over decades, human studies reported varying numbers of rRNA genes in total and on specific acrocentric chromosomes (Stults, Killen et al. 2008). We demonstrate that each individual has a unique number, distribution pattern, and activity of rRNA gene arrays. In other words, everyone has a unique rRNA gene footprint because we inherit one chromosome from each parent, each with its own specific number of genes in the array. Our results are in line with earlier reports demonstrating visible differences in the distribution of rDNA among acrocentric chromosomes in cell lines and primary cells from different individuals (van Sluis, van Vuuren et al. 2020), and presence of inactive NORs in HeLa cells (McStay and Grummt 2008). However, we have shown that the activity of rDNA arrays also varies by chromosome; some arrays are more active, others less active, and some are fully inactive, and the activity status of array is consistent within a specimen. Active arrays are positive for UBF and Treacle and have low methylation at the promoter and 45S gene. In contrast, silent arrays are defined by the absence of UBF and Treacle, high methylation in the promoter and 45S gene, and inaccessible chromatin. Primary cell lines derived from great apes also displayed chromosome-specific array activity, with some inactive arrays, suggesting that this is a common feature of rDNA biology.

The discovery of inactive rDNA arrays allowed us to explore the role of rDNA activity in 3D genome organization, particularly its contribution to the formation of the nucleolar compartment. The nucleolus assembles during interphase around transcriptionally active rRNA genes and contains not only DNA and RNA but also hundreds of proteins (Boisvert, van Koningsbruggen et al. 2007). It exhibits physical cohesiveness and distinct boundaries without having a membrane, but it is surrounded by a pronounced “shell” of heterochromatin (Ferreira, Paolella et al. 1997, Sadoni, Langer et al. 1999, Nemeth and Langst 2011). The T2T human genome assembly revealed that the p-arms of acrocentric chromosomes contain very few coding genes apart from rDNA repeats, but are densely populated with satellite sequences. At the start of each array, rDNA of humans and great apes is flanked by the distal junction (DJ) sequence, a long palindrome that is present on all acrocentric chromosomes and is nearly identical between them. DJ region was proposed to contribute to nucleolar formation by anchoring rDNA to the nucleolar periphery (Floutsakou, Agrawal et al. 2013, Liskovykh, Petrov et al. 2023). DJ sequence was shown associate with the nucleolus in the absence of rDNA in human cells, but in this case it was positioned at the centromere (van Sluis, Gailin et al. 2019). However, centromeres of acrocentric and non-acrocentric chromosomes were also shown to be associated with the nucleolus in various model systems (Haaf and Schmid 1989, Ochs and Press 1992, Carvalho, Pereira et al. 2001). Our analysis of distances to nucleolar boundaries showed that inactive rDNA arrays may not function as NORs as effectively as active arrays, yet they can still associate with nucleoli at rates comparable to the centromeric regions of other chromosomes that do not have rDNA or DJ sequences. Centromeric heterochromatin flanking the active rDNA arrays is naturally associated with nucleoli due to its short distance from the rDNA. When acrocentric chromosomes segregate during cell division, rDNA loci and corresponding centromeres move contiguously into the daughter cells (Potapova, Unruh et al. 2019). Besides, during anaphase, centromeres of all sister chromatids synchronously move together to opposite poles (Maiato and Lince-Faria 2010). We speculate that the physical proximity during sister chromatid segregation in anaphase predisposes centromeres of acrocentric chromosomes with inactive rDNA arrays as well as non-acrocentric centromeres to associate with the nucleolus when it re-assembles upon resumption of transcription, contributing to the formation of the surrounding heterochromatin.

While some reports argue that the rDNA methylation within the coding region does not necessarily inhibit the Pol I transcription (Ghoshal, Majumder et al. 2004, Huang, Zhang et al. 2021), our methyl-sequencing and DNMT1 inhibitor results show that transcriptionally inactive rDNA arrays are silenced by methylation of the promoter and the coding region. DNA methylation, mediated by DNA methyltransferases, is one of the key mechanisms of epigenetic silencing. Inheritance of epigenetic marks was once thought to be impossible due to global demethylation in early embryonic development (Reik, Dean et al. 2001). However, there is evidence that certain genes can escape this demethylation (Lane, Dean et al. 2003), including rRNA genes (Hori, Shimamoto et al. 2021), consistent with our observation that rDNA methylation patterns in HG002 LCLs persisted in iPS stem cells. It has also been shown that methylation marks can be preserved during meiosis, making them heritable across generations (Szyf 2015). In the CHM13 cell line derived from hydatidiform mole where both genome copies are homozygous paternal (Jacobs, Wilson et al. 1980, Fan, Surti et al. 2002), rDNA arrays on both copies of chromosome 22 were silenced, implying the persistence of epigenetic marks from sperm. In the HG002 family trio, the inactive chr.22 rDNA array was inherited from the father, meaning that this epigenetic mark may have been preserved through the germ cell stage, male meiosis, and embryonic development. More investigation will be required to confirm the inheritance of epigenetic features of rDNA arrays.

DNA methylation is not static and can change throughout an organism’s life. It is a key mechanism of epigenetic changes in response to environmental factors (Bonduriansky 2012, Torano, Garcia et al. 2016, Law and Holland 2019, Breton, Landon et al. 2021), which raises a possibility that rDNA silencing can be induced by environmental or other causes, and then propagated. For example, a well-studied case of trans-generational epigenetic inheritance, unrelated to rDNA, is immune priming, where parents can enhance their offspring’s immune defense based on their own immunological experiences, propagating adaptive changes to their progeny (Roth, Beemelmanns et al. 2018). The adaptive value of rDNA silencing is unknown, but there is a possibility that genes containing certain sequence variants in the coding region may be silenced because these variants expressed in rRNA may be disadvantageous. We found that erasing methylation on silent 45S rDNA arrays led to the expression of a specific silent SNP variant in 28S rRNA, but it was unclear if the gene silencing was caused by the presence of this variant. The relationship between 45S and 5S rRNA genes, which too are present as repeat arrays but typically have fewer copies than 45S, is also an open question. Reportedly, there is no correlation between the total numbers of 45S and 5S genes in human individuals (Hall, Turner et al. 2021), but the information on the physiological lower and upper limits of both types of rRNA genes required for viability is lacking. It is tempting to speculate that epigenetic modifications may be needed to maintain a sustainable ratio of active 45S/5S genes for a balanced rRNA output that can sustain ribosome biogenesis. Our findings demonstrate the existence and potential heritability of rDNA epigenetic patterns, but their underlying causes and adaptive significance remain to be understood.

## Acknowledgments

We are thankful to the Sequencing and Discovery Genomics, Flow Cytometry, Tissue Culture and Microscopy core facilities at the Stowers Institute for enabling many of our experiments. We are grateful to Kateryna Makova and Barbara McGrath from Pennsylvania State University for providing primate cell lines. We thank members of Gerton lab for discussions. We appreciate Mark Miller’s help with illustrations. This work was supported by funding from Stowers Institute and by the Intramural Research Program of the National Human Genome Research Institute, US National Institutes of Health. This work utilized the computational resources of the NIH HPC Biowulf cluster (https://hpc.nih.gov).

## Materials and Methods

### Cell culture

Human lymphoblastoid cell lines (LCL) including HG02055, HG02054, HG01109, HG01243, HG03492, HG02723, HG02082, HG02053, HG002 (GM24385), HG003 (GM24149), HG004 (GM24143), and induced pluripotent stem cells (iPS) GM26105 and GM27730 were obtained from Coriell. All LCL cell lines were cultured in RPMI 1640 (Gibco) with L-glutamine supplemented with 15% fetal bovine serum (FBS). iPS cells were initially cultured on MEF feeder cell monolayer in the DMEM:F12 medium containing 20% KnockOut Serum Replacement (Gibco), 0.1mM non-essential amino acids (Gibco), 55µM β-mercaptoethanol, 10ng/ml bFGF (Corning) and 10 µM ROCK inhibitor Y-27632 (STEMCELL Technologies), and then adapted to the feeder-free culture. For iPS adaptation to feeder-free culture colonies were seeded on Matrigel-coated plates in mTeSR Plus medium (STEMCELL Technologies) and passaged at least three times until no feeder cells were observed. CHM13 cells were obtained from a case of a complete hydatidiform mole at Magee-Womens Hospital (Pittsburgh) as part of a research study IRB MWH-20-054, and immortalized using human telomerase reverse transcriptase (hTERT) to develop an established cell line. After initial culture in complete AmnioMax C-100 Basal Medium (Thermo Fisher Scientific) cells were propagated in DMEM:F12 supplemented with 1x non-essential amino acids (Gibco), 1x Insulin-Transferrin-Selenium (Gibco), 1mM sodium pyruvate (Gibco), and 10% FBS. All primate cell lines were obtained from Makova lab (Penn State University). Fibroblasts from Bonobo, Bornean orangutan, and gorilla were cultured in Alpha MEM with L-glutamine (Gibco) supplemented with 10% FBS. Fibroblasts from Sumatran orangutan were grown in Alpha MEM with L-glutamine supplemented with 15% FBS and 1x non-essential amino acids (Gibco). Chimpanzee LCL cell line was grown in RPMI 1640 (Gibco) supplemented with 10% FBS, 1x sodium pyruvate (Gibco), and 1x non-essential amino acids (Gibco). All cell lines were grown in a 37°C incubator with 5% CO_2_.

### Chromosome spreads, Fluorescent In-Situ Hybridization (FISH), and immuno-FISH

For the preparation of chromosome spreads, cells were blocked in mitosis by the addition of Karyomax colcemid solution (0.1 µg/ml, Life Technologies) for 6-7h. Adherent fibroblast cells were collected by trypsinization. Collected cells were incubated in hypotonic 0.4% KCl solution for 12 min and prefixed by addition of methanol:acetic acid (3:1) fixative solution (1% total volume). Pre-fixed cells were spun down and then fixed in Methanol:Acetic acid (3:1). Spreads were dropped on a glass slide and incubated at 65°C overnight. Before hybridization, slides were treated with 0.1mg/ml RNAse A (Qiagen) in 2xSSC for 45 minutes at 37°C and dehydrated in a 70%, 80%, and 100% ethanol series for 2 minutes each. Slides were denatured in 70% deionized formamide/2X SSC solution pre-heated to 72°C for 1.5 min. Denaturation was stopped by immersing slides in 70%, 80%, and 100% ethanol series chilled to -20°C. Labeled DNA probes were denatured separately in a hybridization buffer by heating to 80°C for 10 minutes before applying to denatured slides. Specimens were hybridized to the probes under a glass coverslip or HybriSlip hybridization cover (GRACE Biolabs) sealed with the rubber cement or Cytobond (SciGene) in a humidified chamber at 37°C for 48-72hours. After hybridization, slides were washed in 50% formamide/2X SSC 3 times for 5 minutes per wash at 45°C, then in 1x SSC solution at 45°C for 5 minutes twice and at room temperature once. For biotin detection, slides were incubated with streptavidin conjugated to Cy5 (Thermo) for 2-3 hours in PBS containing 0.1% Triton X-100 and 5% bovine serum albumin (BSA), and then washed 3 times for 5 minutes with PBS/0.1% Triton X-100. For immuno-FISH, slides labeled by FISH were subjected to antigen unmasking in hot (65°C) Citrate buffer, pH 6.0, for 1 hour before processing for immunofluorescence. Slides were blocked with 5% bovine serum albumin (BSA) in PBS/0.1% Triton X-100. Primary antibody (rabbit polyclonal anti-UBF, Novus Biologicals, cat.# NBP1-82545) and secondary antibody (goat anti-rabbit Alexa Fluor 647, Thermo) were diluted in 2.5% (weight/volume) BSA/PBS/0.1% Triton X-100. Specimens were incubated with primary antibody at a minimum overnight, washed 3 times for 5 minutes, incubated with secondary antibody for 2-4 hours and washed again 3 times for 5 minutes. All washes were performed with PBS/0.1% Triton X-100. Slides were mounted in Vectashield containing DAPI (Vector Laboratories). Wide-field images were acquired on the Nikon TiE microscope equipped with 100x objective NA 1.4 and Prime 95B sCMOS camera (Photometrics). Z-stack images were acquired on LSM780 confocal microscope (Zeiss) using the 63x/1.40 NA oil objective, or the Nikon TiE microscope equipped with 100x objective NA 1.45, Yokogawa CSU-W1 spinning disk, and Flash 4.0 sCMOS camera (Hamamatsu).

### Estimating rDNA copy number and activity from FISH images

Image processing was performed in FIJI and Python. For chromosome identification in spreads from human cell lines, labeling centromeric satellite 14/22 and PML locus on the q-arm of chromosome 15 was sufficient to identify all rDNA-containing chromosomes. Primate rDNA-containing chromosomes were identified as homo sapiens (hsa) homologs based on the labeling with human chromosome paints and morphological features. For chimpanzee and bonobo, painting hsa 14 and hsa 21 was sufficient to identify all rDNA-containing chromosomes, and for gorilla, hsa 22 paint alone was sufficient. Chromosome Y was identified by morphology. For Sumatran and Bornean orangutans, all rDNA-containing chromosomes were painted on separate slides, and these data were aggregated across all slides.

For manual image quantifications, performed for chromosome spreads from human CHM13 cells, chimpanzee, bonobo and gorilla cells, sum intensity projections of confocal Z-planes were generated, and individual rDNA arrays were segmented based on threshold applied to the entire image. The fluorescence intensity of the regions of the same chromosomes that did not contain the rDNA was used to subtract the local background. The background-subtracted integrated intensity was measured for each array. For semi-automated quantification performed for chromosome spreads from human HPRC panel, Sumatran orangutan and Bornean orangutan, wide-field single Z-plane images were used. rDNA-containing acrocentric chromosomes were segmented using a Cellpose model trained on 2-channel images including the DAPI and rDNA signals. rDNA regions were also segmented using a trained Cellpose model. The chromosome segmentations were examined and, if necessary, curated manually in Napari. rDNA and UBF intensities for each array were measured after subtracting the fluorescence background for the respective chromosomes, and the fraction of the total rDNA and UBF fluorescence intensity in the cell was calculated for each array.

The sum of all intensities of all rDNA loci represented the total amount of rDNA per cell. The total rDNA copy number was estimated from Illumina sequencing data (see “Estimating rDNA copy number from k-mer coverage”). The fraction of the total rDNA fluorescence intensity was used as a proportion of the total rDNA copy number to determine the number of rDNA copies on specific chromosomes in each chromosome spread.

### Estimating rDNA copy number from k-mer coverage in HPRC samples

Ribosomal DNA copy numbers were estimated from k-mer frequencies in the Illumina PCR-free short read whole genome sequencing data. The data source for the HPRC samples (Fig. 1C) was the 1000 Genomes Project phase 3 samples which are available on the Sequence Read Archive, and the source for the HG002 trio was a study from (Gunjan Baid 2020). NCBI entry KY962518.1 https://www.ncbi.nlm.nih.gov/nuccore/KY962518.1 was used as a reference sequence for human 45S rDNA. The 18S copy number served as a proxy for the greater 45S unit, as each unit contains a single 18S segment. A custom pipeline counted k-mers of size 31 from the 18S consensus in short read Illumina sequencing data and normalized it to counts of 31mers from G/C matched windows elsewhere in the rDNA containing chromosomes. The matched windows were of similar size to the 18S, and ten of these were randomly selected per rDNA-containing chromosome. Any k-mers which also occurred outside the matched windows were removed to ensure that counts were exclusively from the matched windows. k-mer sets were filtered to remove those with whole genome sequencing counts greater than three standard deviations from the mean of the set, or those which were missing entirely. Counts were divided by their genomic multiplicity. Finally, the median count from the 18S k-mers was divided by the median count of the matched windows to yield a copy number approximation. A pipeline referred to as CONKORD (version 7) was used for this process, except for HG002 trio for which very similar version 6 was used.

### Quantification of nucleolar associations from immuno-FISH images

CHM13 cells were grown on #1.5 glass coverslips, fixed in 4% paraformaldehyde in PBS for 15 minutes, and permeabilized with 0.1% Triton X-100 in PBS and stored in 25% glycerol/PBS at 4°C. Before hybridization, coverslips were subjected to two freeze-thaw cycles by dipping into liquid nitrogen, treated with 0.1 N HCl for 5 min, washed twice in 2× SSC buffer, and pre-incubated in 50% formamide/2× SSC overnight. Fluorescently labeled probes were pre-denatured for 2 minutes at 85°C, followed by incubation with the specimen for 5 min at 85°C, and hybridized under HybriSlip hybridization cover (GRACE Biolabs) sealed with Cytobond (SciGene) in a humidified chamber at 37°C for 48-72 hours. After hybridization, slides were washed in 50% formamide/2X SSC 3 times for 5 minutes per wash at 45°C, then in 1x SSC solution at 45°C for 5 minutes twice and at room temperature once. Slides were washed again in 0.1% Triton X-100 in PBS and blocked with 5% BSA in PBS/0.5% Triton X-100. Primary and secondary antibodies were diluted in 5%BSA/PBS/0.1% Triton X-100. Specimens were incubated with primary antibody overnight, washed 3 times for 5 minutes, incubated with secondary antibody for several hours, and washed again 3 times for 5 minutes. All washes were performed with PBS/0.1% Triton X-100. Vectashield containing DAPI was used for mounting.

Z-stack confocal images were acquired using a Nikon TiE microscope equipped with a 60x objective lens NA 1.4. Since interphase CHM13 nuclei are flat, distances to the nearest nucleolar boundaries were measured on maximum intensity projections. Nuclei were segmented based on the thresholded DAPI channel and converted into nuclear masks. Polyploid or otherwise abnormal nuclei were excluded from the analysis. Nucleoli were segmented using the thresholded nucleolin channel and converted into masks, that were then transformed into Euclidean Distance Transform (EDT) maps, where pixel values represent the distance from the nearest boundary of the mask. This transformation was performed using the ‘test sedt jru v1’ plugin for ImageJ/FIJI available at http://research.stowers.org/imagejplugins. The EDT output image was inverted by multiplying it by -1 to produce positive distance values. Chromosome markers were segmented based on threshold and size criteria, and the points outside nuclear masks were excluded. For a random points control, two single-pixel points were placed in each nuclear mask using the ‘random’ function built into ImageJ. The values of the EDT output image, that indicate the closest distances in pixels, were measured at the positions of segmented marker foci in each nucleus. These distances were converted to microns using the known scale of the objective lens. For foci inside the nucleolar masks, negative distance values were converted to zero. The threshold for nucleolar association was set at 0.5 µm from the nearest nucleolar boundary.

### Chromosome isolation, sorting, and methyl-seg analysis

To induce mitotic arrest, CHM13 cells were treated with 100µM Monastrol (Tocris) and 10µM pro-Tame (R&D Systems) for 7-10 hours. Mitotic cells were collected by shake-off. Collected cells were treated with 0.1 µg/ml Colcemid for 15 min, and incubated in hypotonic swelling buffer containing 45mM KCl, 10 mM MgSO4, 0.5mM Spermidine and 0.2mM Spermine for 12 min and collected by centrifugation at 335g for 5min. Cells were lysed in 1ml polyamine buffer (150 mM Tris pH 7.5, 80 mM KCl, 2 mM EDTA, 0.5 mM EGTA, 3 mM DTT, 0.25% Triton X-100, 0.2 mM spermine, 0.5 mM spermidine) on ice for 30 minutes with periodic vortexing. Chromosomes were labeled with overnight at 4°C with 5 µg/ml Hoechst 33258 and 50 µg/ml Chromomycin A3 in the presence of 10 mM MgSO4, vortexed and passed through an 8 µm filter. Two or more hours before analysis, 10 mM sodium citrate and 25 mM sodium sulfite were added to the suspension. Chromosome suspensions were analyzed and sorted on a BD Bioscience FACSymphony S6 Cell Sorter with a 70 µm tip at 70 psi using 1x PBS, diluted from 10x Preservation Free ClearSort Sheath Fluid (Leinco Technologies Cat# S632), containing 0.1% of a non-ionic surfactant, Poloxamer 188 (Mirus Cat#6230A). The sample pressure differential was kept as low as possible to maintain a tight sample core. Chromomycin A3 was excited with a 445nm OBIS laser at 75 mW (Coherent); emission wavelengths were collected centered at 510nm. Hoechst 33258 was excited with a Genesis G4 355 UV laser at 100 mW (Coherent); emission wavelengths were collected centered at 450nm. Threshold gating was performed using the Hoechst 450nm emission detector to remove debris.

The target chromosome populations were sorted with a 2-pass strategy. The result of the first pass (sort) yielded samples with purities between 70-80%. These sorted samples were re-sorted yielding target population purities >95%. Flow cytometry data were collected and analyzed with FACSDiva (BD Bioscience) and FlowJo software (BD Bioscience).

Genomic DNA from sorted chromosomes was isolated using QIAamp DNA Micro Kit (Qiagen) according to the manufacturer’s protocol. Methyl-seq libraries were generated from 2.4-6.6ng of DNA isolated from sorted chromosomes, as assessed using the Qubit Fluorometer (Life Technologies). Libraries were made according to the manufacturer’s directions for the NEBNext Enzymatic Methyl-seq kit (New England Biolabs, Cat. No. E7120S) using the S220 Focused-ultrasonicator (Covaris) to shear to 300bp and 9 cycles of library PCR amplification. Resulting short fragment libraries were checked for quality and quantity using the Bioanalyzer (Agilent) and Qubit Fluorometer (Life Technologies). Libraries were pooled, requantified and sequenced as 150bp paired reads on a mid-output flow cell using the Illumina NextSeq 500 instrument, utilizing RTA and instrument software versions current at the time of processing. Paired-end methyl sequencing reads were trimmed with trim galore using the default settings of paired-end mode. Reads were then aligned to chm13 (v2.0) using the bisulfite sequencing aligner Bismark (v0.24.0). The default end-to-end alignment mode was used. Alignments were deduplicated and extracted into bed coverage format to give a C and T count at each reference position. The purity of the flow sorted chromosomes was assessed by finding the fraction of aligned reads with a best hit to the chromosome we attempted to isolate over the total number of aligned reads. Acrocentrics were split into 10 kb bins, and the average percent methylation for each bin was found by taking the total number of Cs divided by total coverage. CpG sites were not included in this calculation if they were covered by less than five reads. To assess methylation at the rDNA alone, the above process was repeated except the rDNA reference KY962518.1 was used in place of chm13 genome. Methylation percentages were plotted for each CpG site that was covered by at least 10 reads.

### Whole genome Methyl-seq analysis

Genomic DNA from untreated and 5uM GSK-3484862 treated cells was isolated using QIAamp DNA Micro Kit (Qiagen). A Methyl-seq library was generated from 200ng of DNA, as assessed using the Qubit Fluorometer (Life Technologies). The library was made according to the manufacturer’s directions for the NEBNext Enzymatic Methyl-seq kit (New England Biolabs, Cat. No. E7120S) using the S220 Focused-ultrasonicator (Covaris) to shear to 300bp and 4 cycles of library PCR amplification. The resulting short fragment library was checked for quality and quantity using the Bioanalyzer (Agilent) and Qubit Fluorometer (Life Technologies). The library was sequenced as 150bp paired reads on a high-output flow cell using the Illumina NextSeq 500 instrument, utilizing RTA and instrument software versions current at the time of processing. Bismark was used to align reads to the rDNA reference KY962518.1 using the local parameter to allow soft-clipping. Reads were not trimmed prior to analysis. The bismark deduplicate and extract methylation commands were used to generate bed coverage files with C and T counts for each CpG site in the rDNA. Sites were filtered if they were covered by less than 10 reads.

### Fiber-Seq Analysis

Publicly available Fiber-sequencing data from the CHM13 cell line was obtained from the NCBI Sequence Read Archive (SRA) under accession number SRR16356599. The fibertools software (https://github.com/fiberseq) was used for analysis. Circular consensus sequences were generated from subreads with the option to keep kinetics. M6A methylation was then predicted using ft m6a-predict on the circular consensus sequence. To generate an aligned file, the m6a-predicted bam file was first converted to a fastq format using samtools fastq, while transfering all tags to the header. The fastq data was then aligned to NCBIs KY962518.1 rDNA reference using minimap with parameters optimized for PacBio. The alignment was performed with multi-threading and the resulting alignments were saved in SAM file format. The SAM file was then converted to BAM format using samtools view before sorting. Reads were filtered to ensure that they fully covered the 28S region of the reference. Individual alignments were separated into new bam files, and these files were sorted based on the presence or absence of an A to G SNP in the 28S rDNA (position 11253 in the KY962518.1 reference). Samtools mpileup was used to find the sequence at this position. The tool modkit was used to translate the Mm and Ml tags for every alignment into bedmethyl files containing m6a percentages. The command modkit pileup was run with the --force-allow-implicit and --motif A 0 parameters set.

### DNA, RNA sequencing and variant calling analysis

Publicly available DNA sequencing data from the CHM13 cell line was obtained from the NCBI Sequence Read Archive (SRA) under accession number SRX1009644. CHM13 RNA was isolated in-house using the NEB Monarch Total RNA Miniprep Kit following the product specifications. Total RNA was further purified with DNAseI clean up. TruSeq Stranded Preparation kit was used to prepare libraries according to manufacturer’s instructions using TruSeq Universal Adaptors. The sequencing was performed on an Illumina NextSeq 500 (mid output) with strand-specific, paired-end 2×75 base reads. The RNA-seq data was trimmed using Trim Galore (v0.6.7) to remove adaptors and bases with a Phred score less than 20. The trimmed reads were then aligned to the KY962518.1 NCBI rDNA reference sequence using BWA MEM (v0.7.17). Aligned SAM files were converted to BAM format, sorted, and indexed using samtools (v1.18). Variant calling was performed by LoFreq. VCF files were filtered to remove variants with a quality score less than 30. To authenticate the variants called by LoFreq, a summary file was generated using Samtools mpileup. This mpileup file provided a detailed base-pair resolution of the read alignments, which was used to cross-check the presence and frequency of the identified variants. Downstream analysis involved examining the mpileup summary file to ensure that the variants had consistent support across multiple reads and were not artifacts of sequencing or alignment errors.

### CHM13 rRNA gene methylation analysis from ONT reads

Oxford Nanopore Technology (ONT) reads were basecalled with modifications using Guppy v6.5.7 with the following command:

guppy_basecaller --num_callers 1 --cpu_threads_per_caller $cpus --compress_fastq -- do_read_splitting -i $input_path -s $output_path -c dna_r9.4.1_450bps_modbases_5mc_cg_sup_prom.cfg -x “cuda:all” -r Reads were then aligned to CHM13 using winnowmap v2.03 (Jain, Rhie et al. 2022) and samtools v1.19 using the following command:

winnowmap -t $cpus -W chm13.repetitive_k15.txt --eqx --MD -a -y -x map-ont -I 12g chm13v2.0.fa

$reads_fq | samtools view -@$cpus -h -O SAM -F 260 | samtools sort -@$cpus -O BAM --write-index - o $out.bam##idx##$out.bam.bai

Reads were assigned to chromosomes using a custom python script, linked in the GitHub. For reads with multiple alignments, if the best alignment contained 1.5x more matching bases than the next best alignment, it was assigned as specific to that chromosome. Otherwise, reads were marked unassignable and left out of further analyses. From there, the chromosome-split ONT reads were aligned to the rotated reference unit of the rDNA array KY962518.1 using minimap2 (Li 2018). Unmapped reads were filtered, as were reads not meeting 90% alignment block and 90% identity, and suspected chimeric reads containing inverted units were removed. Aggregated methylation percentages at all CpGs were obtained using modkit v0.3.1 (https://github.com/nanoporetech/modkit) with the following command:

modkit pileup --threads $cpus --ref {KY962518-ROT. --cpg --combine-strands --ignore h --log $out.log

$out_dir/$chr.bam $out_dir/$chr.modkit.bed

Finally, the aggregated methylation results were visualized for each acrocentric chromosome, with the average signal computed for non-overlapping 200 bp windows across the entire reference sequence.

### Probes and antibodies used in this study

Centromere probe for chr.13 was generated in-house by PCR-amplification of CHM13 genomic DNA using the following primers:

Chr13_Fwd:5’-GGGAATTCAAATAAAAGGTAG-3’ ; Chr13_Rev: 5’-CCAAATGTCCACATCCAGA-3’

Amplicon sequence: GGGAATTCAAATAAAAGGTAGacagcagcattctcagaaatttctttctgatgtctgcattcaactcatagagttgaagattccctttcatagagcaggtttgaaacactctttctggagtaTCTGGATGTGGACATTTGG .

The PCR amplicon was labeled with Biotin-16-dUTP using the nick translation kit (Enzo Life Sciences) and detected with streptavidin conjugated to Cy5 (Thermo)

The oligonucleotide biotin-labeled probe for WaluSat was from IDT:

5’-/5Biosg/AGA AAG GGA TAG GAG TGA AGA ACA CAG GTC GCT GCA TTT AGA AAG GAG GCG GGG TCA GAG GAA T /3Bio/ -3’

| Commercially available reagents: |  |  |
| --- | --- | --- |
| Probe | Cat.# | Source: |
| CenSat D14Z1/D22Z1 | LPE 014 | Cytocell |
| CenSat D15Z4 | LPE 015 | Cytocell |
| CenSat D7Z1 | LPE 07 | Cytocell |
| CenSat D8Z2 | LPE 08 | Cytocell |
| CenSat D12Z3 | LPE 012 | Cytocell |
| CenSat D20Z1 | LPE 020 | Cytocell |
| CenSat DXZ1 | LPE 0X | Cytocell |
| Chr.22 qter | LPT 22Q | Cytocell |
| Human 45S rDNA probe | BAC RP11-450E20 | Empire Genomics |
| Human 5S rDNA probe | WI2 1774C11, WI2 2801N10 | Empire Genomics |
| Chr.14 near-cen marker | RP11-55G7 | Empire Genomics |
| Chr.21 near-cen marker | BAC RP11-846C20 | Empire Genomics |
| Chr.22 near-cen marker | BAC RP11-806D22 | Empire Genomics |
| Antibody |  |  |
| Anti-UBF | NBP1-82545 | Novus Biologicals |
| Anti-Nucleolin | ab70493 | Abcam |
| Anti-Treacle/TCOF1 | ab224544 | Abcam |

Secondary antibodies for immunofluorescence (AlexaFluor 647 conjugates) were obtained from Life Technologies and used at 1:500 dilution.

