## Supplementary figures 1-7 for "Epigenetic control and inheritance of rDNA arrays"

### Supplementary Figure 1. rDNA copy number quantifications for HPRC panel

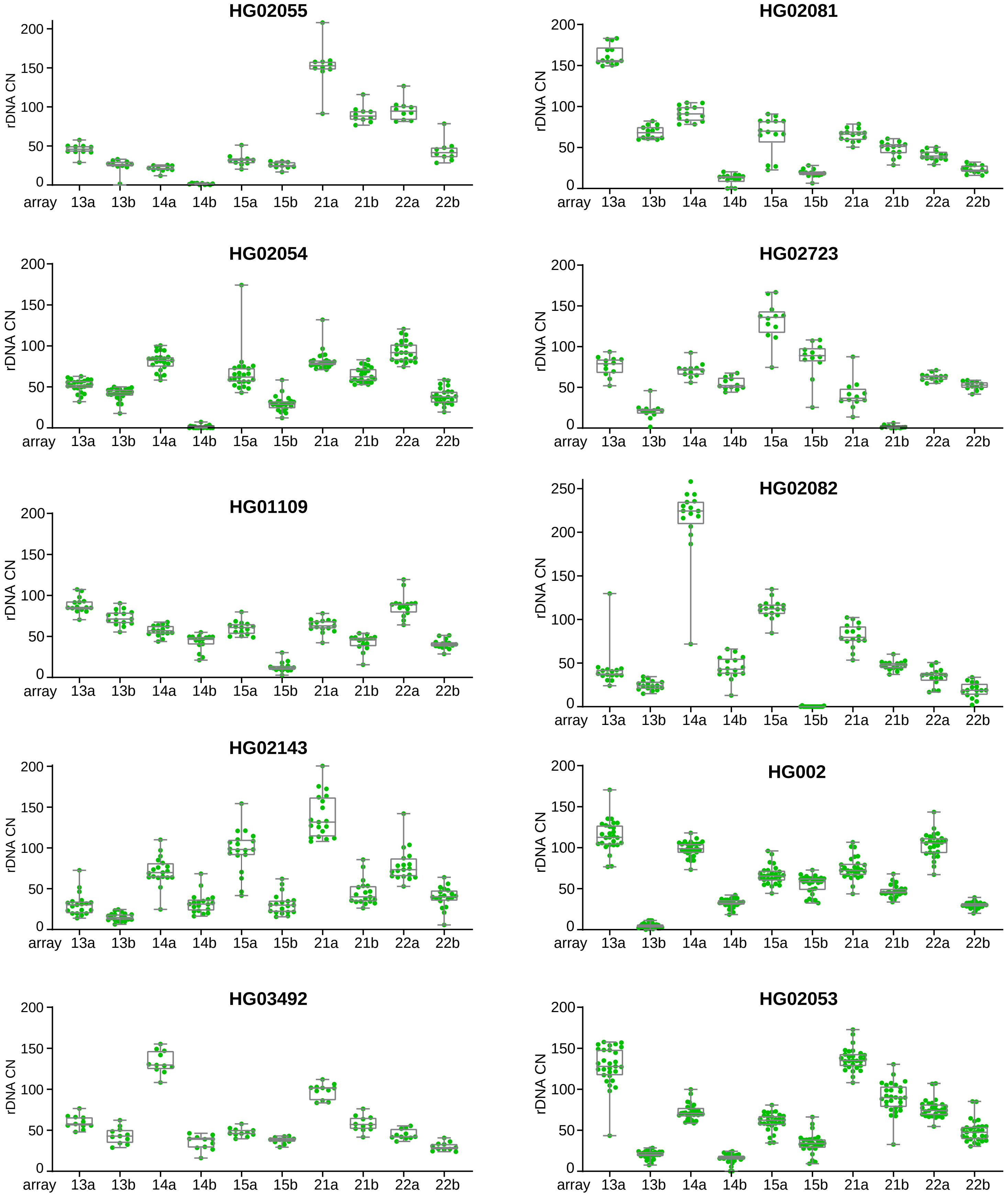


Quantification of rDNA copy number on acrocentric arrays in selected panel of HPRC cell lines shown in Figure 1B. The boxes represent the interquartile range, with the edges indicating the upper and lower quartiles. The line inside the boxes indicates the median. Whiskers show the range from minimum to maximum values. All individual data points are shown.

#### Supplementary Figure 2. rDNA array size and activity quantifications for HPRC panel

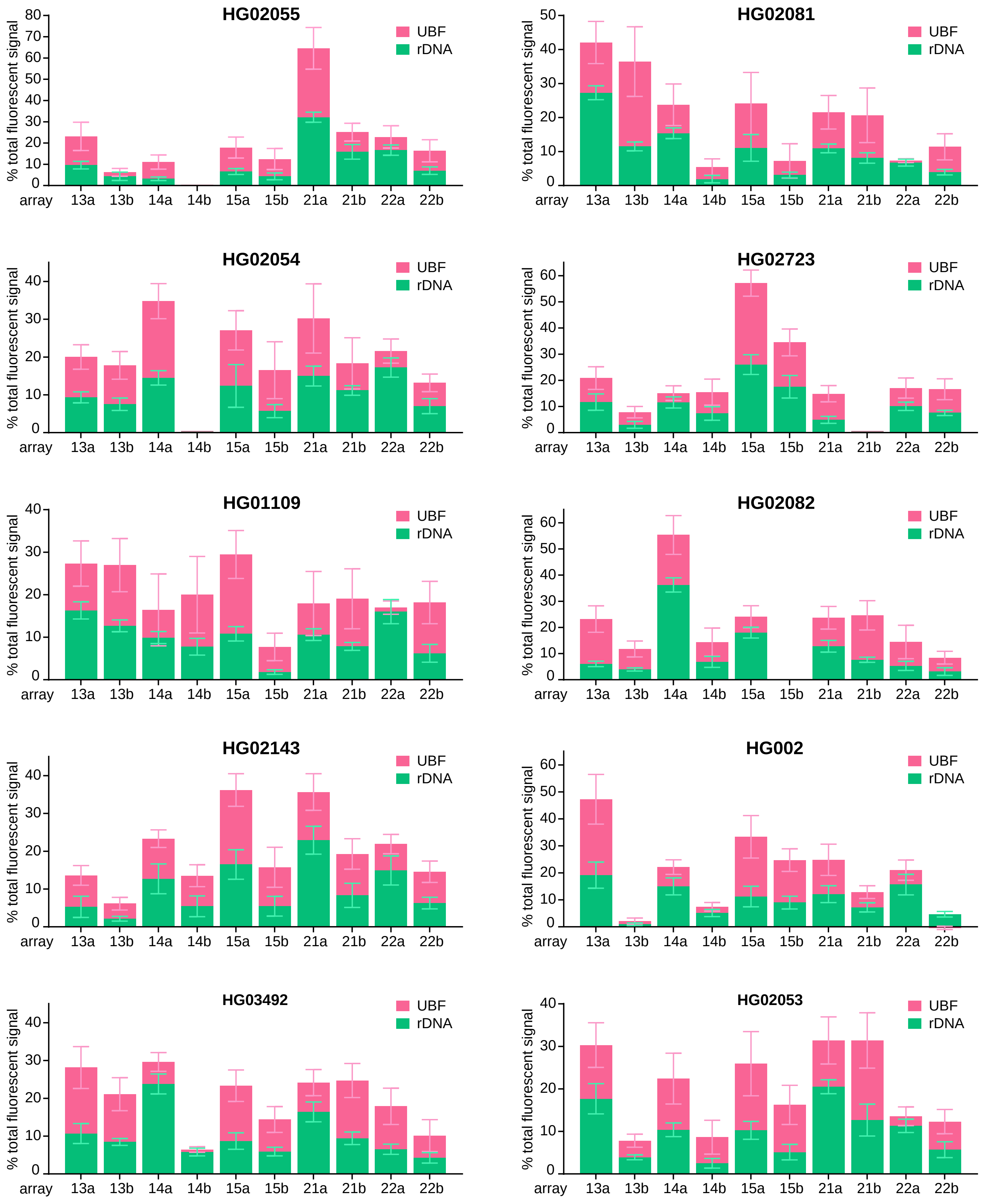


Quantification of rDNA FISH and UBF antibody labeling on acrocentric arrays in selected panel of HPRC cell lines shown in Figure 1C and 1D. rDNA FISH and UBF antibody signals were measured as fractions of the total fluorescent intensity in the chromosome spread. The green and pink sections of the bars represent averages of rDNA and UBF, respectively. Error bars denote standard deviation.

#### Supplementary Figure 3. Activity of rDNA arrays in great apes

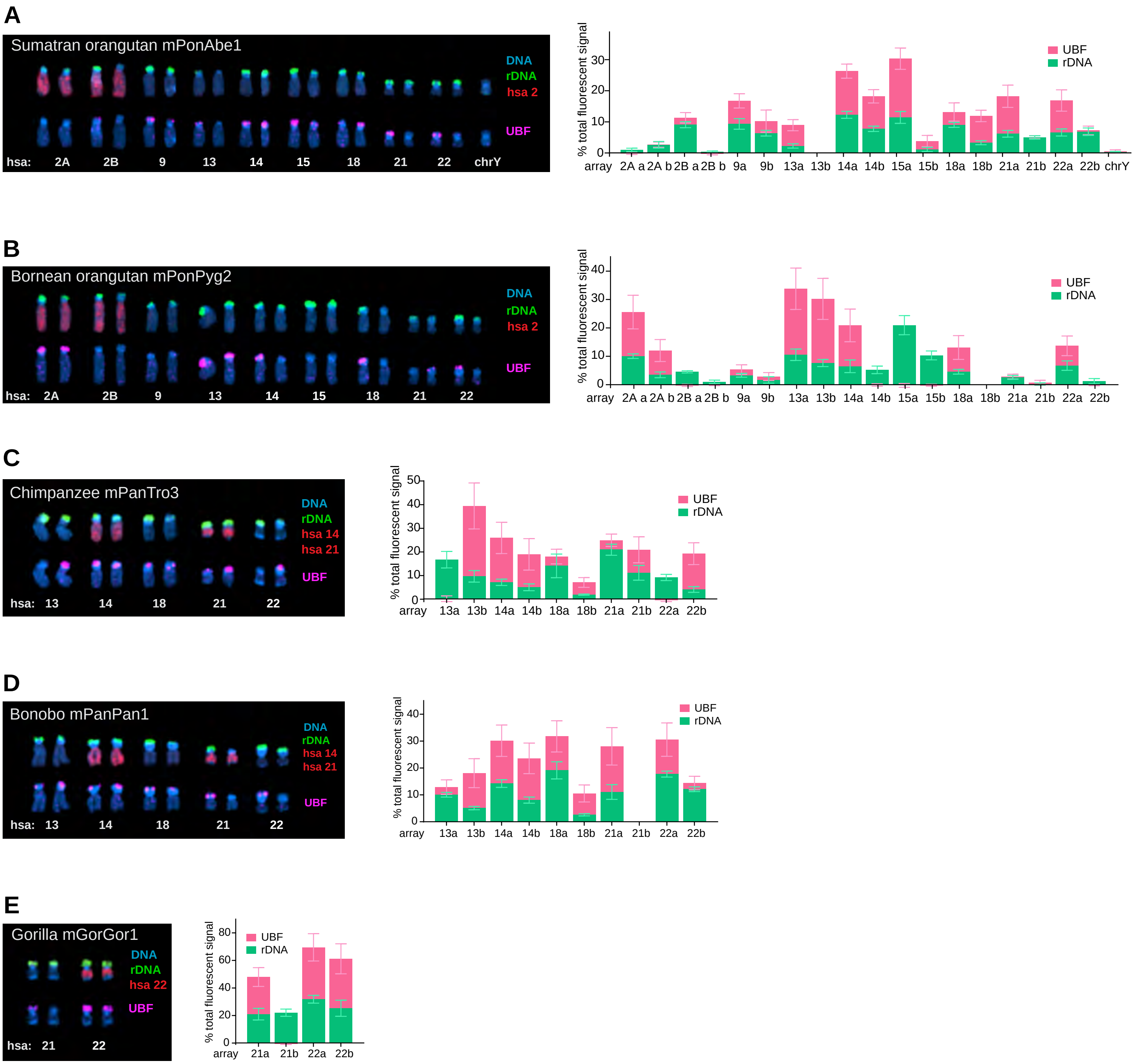


Representative karyograms of rDNA-bearing chromosomes in cells from Sumatran orangutan(A), Bornean orangutan (B) chimpanzee (C), bonobo (D) and western lowland gorilla (E). Chromosomes were identified as homo sapiens (hsa) homologues. The top rows of chromosomes show FISH labeling with rDNA probe (green) and indicated human whole chromosome paints (red). The bottom rows show labeling with UBF antibody (magenta). DNA was counter-stained with DAPI. Corresponding quantifications of rDNA FISH and UBF antibody labeling are shown on the right. The green and pink sections of the bars represent average intensities of rDNA and UBF, expressed as percentage of total signal. Error bars denote standard deviation.

#### Supplementary Figure 4. Validation of rDNA-adjacent chromosome-specific markers and histograms of distances to the nearest nucleolar boundary

**A**

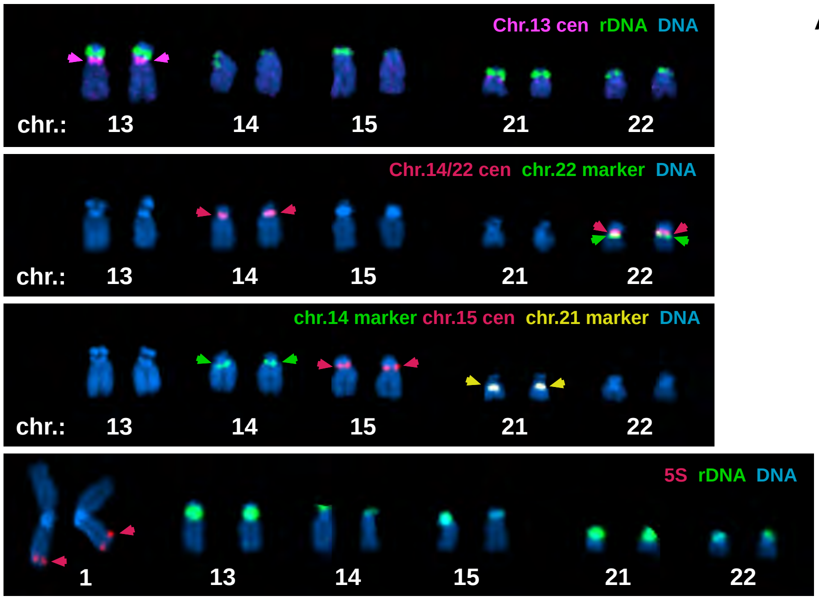


**A.** Acrocentric karyograms from CHM13 cells labeled by FISH with indicated BAC probes used in Figure 2 E-F to confirm localization.
First panel: 45S rDNA RP-11 450E20 (green) and in house-generated probe for centromere 13 (magenta, detailed in Materials and Methods). Second panel: centromere probe D14Z1/ D22Z1 alpha-satellite (red) and chromosome 22 near-centromere marker RP-11 806D22 (green). Third panel: chromosome 14 near-centromere marker RP-11 55G7 (green), chromosome 15 centromere probe D15Z4 (red) and chromosome 21 near-centromere RP-11 846C20 (yellow). Fourth panel: 5S rDNA array probe – combination of WI2 1774C11 & WI2 280N10 (red), and 45S rDNA RP-11 450E20
(green). DNA was counter-stained with DAPI.

**B**

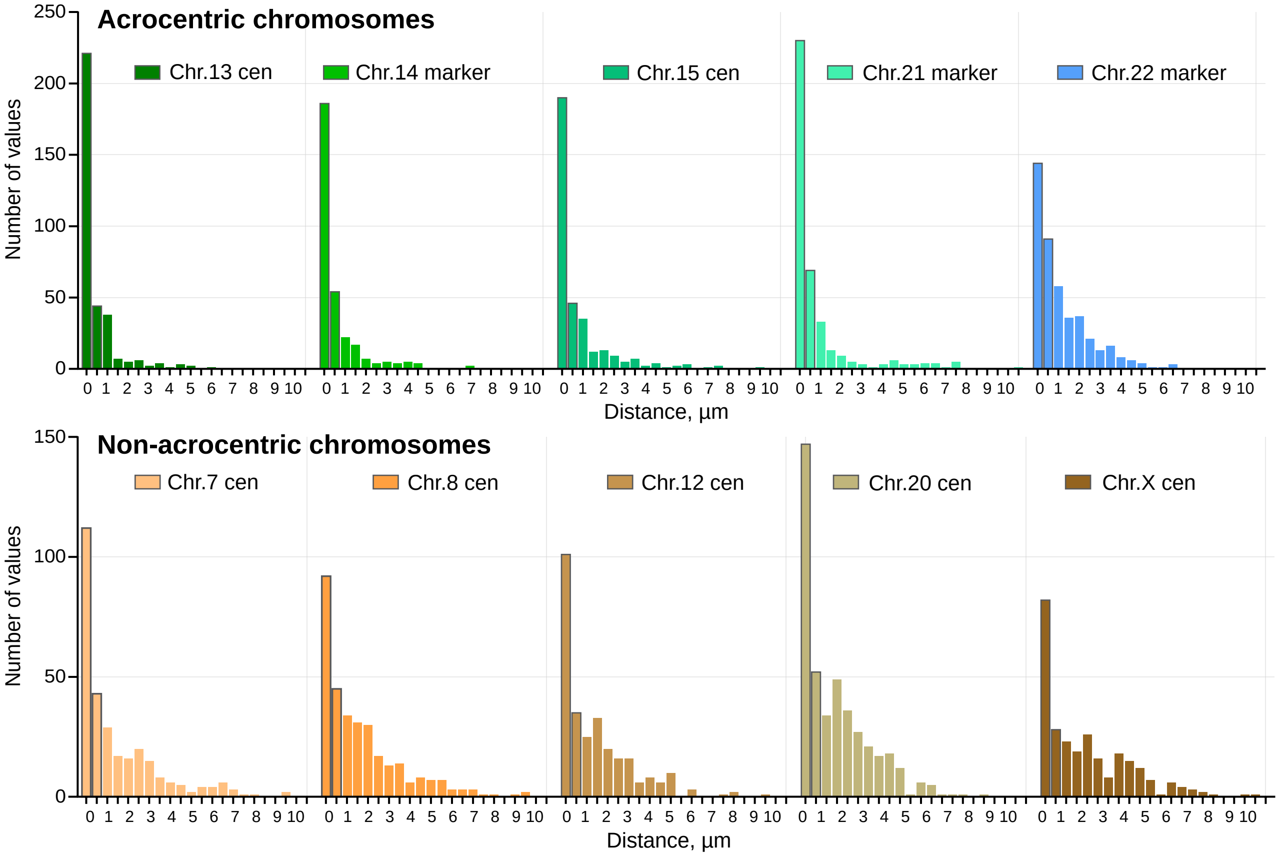


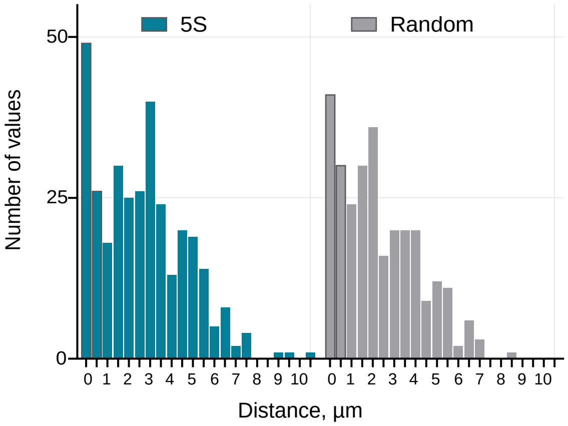


**B.** Distributions of distances from the nucleolar boundary for indicated chromosome markers.
First row – acrocentric chromosomes, second row – non-acrocentric chromosomes, third row – 5S rDNA and the Random points control. First two bars (outlined) represent nucleolar-associated values less than 0.5µm away from the nearest nucleolar boundary.

#### Sup. Figure 5. Methylation of rRNA genes in sorted chromosomes from CHM13 cell line

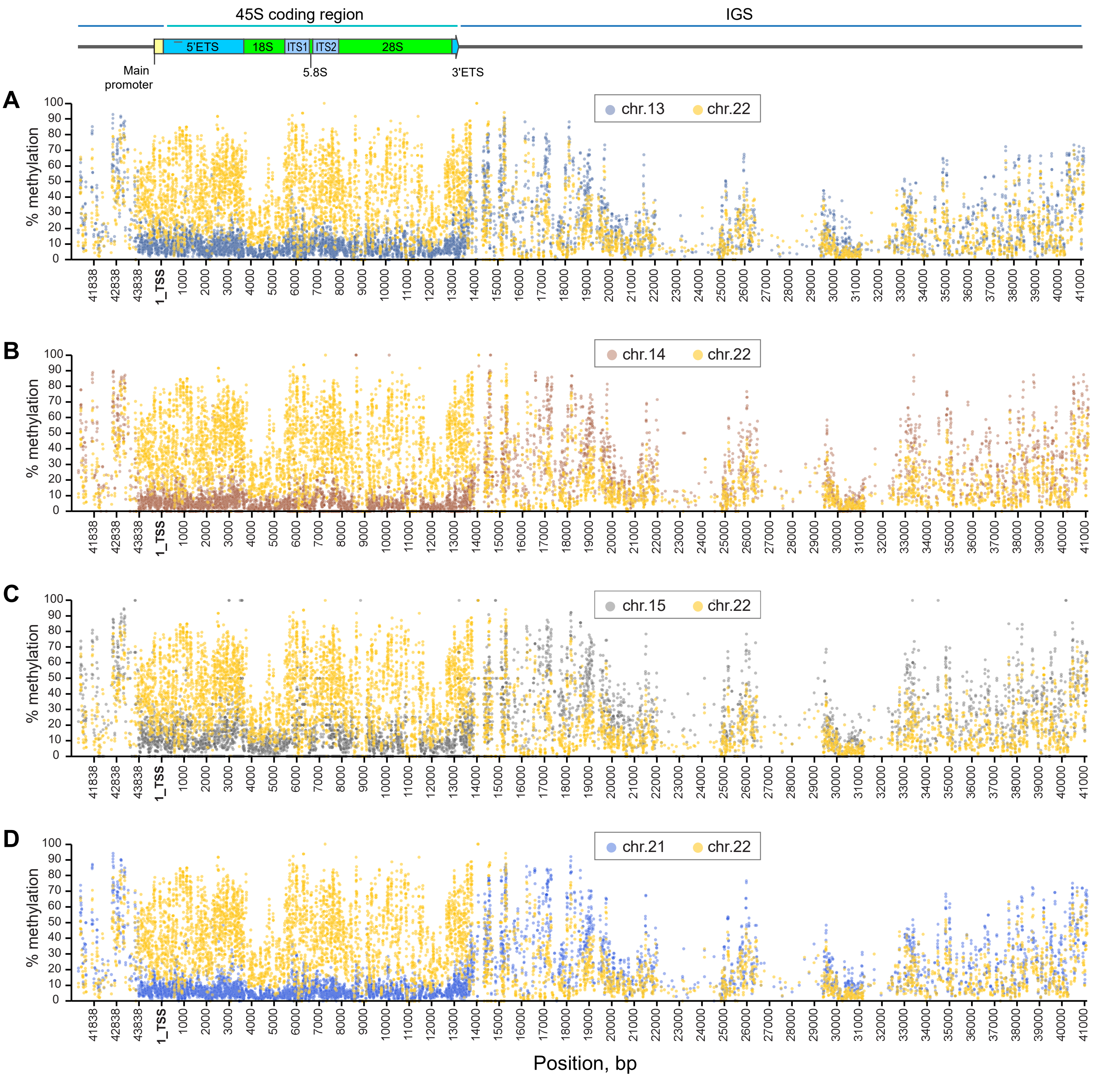


Short-read methyl-sequencing analysis showing the percent methylation of each cytosine base in rDNA reads across the gene. Individual plots display rDNA methylation for reads mapped to chromosome 13 (A), chromosome 14 (B), chromosome 15 (C), and chromosome 21 (D). Chromosome 22 rDNA reads are indicated by yellow circles.

#### Sup. Figure 6. Related to rDNA methylation analysis in CHM13 cell line

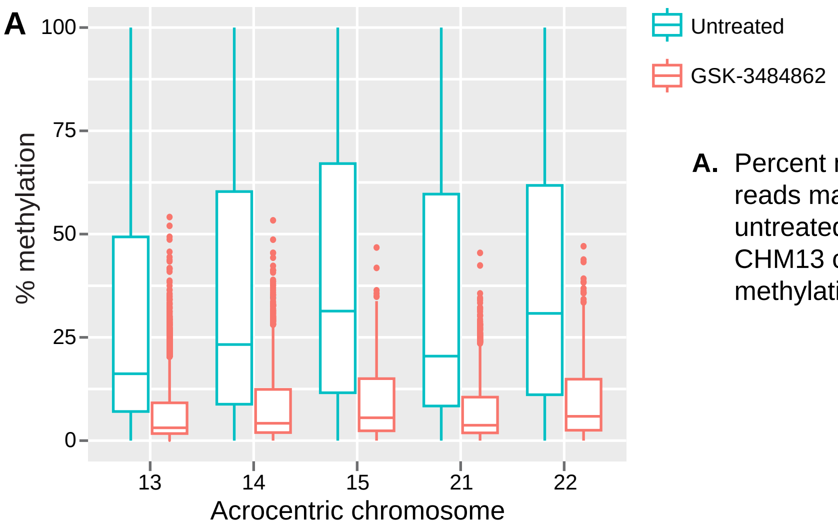


**A.** Percent methylation of non-rDNA methyl-sequencing reads mapped to all acrocentric chromosomes in untreated and DNMT1 inhibitor GSK-3484862 treated CHM13 cells. The DNMT1 inhibitor reduces DNA methylation levels genome-wide.

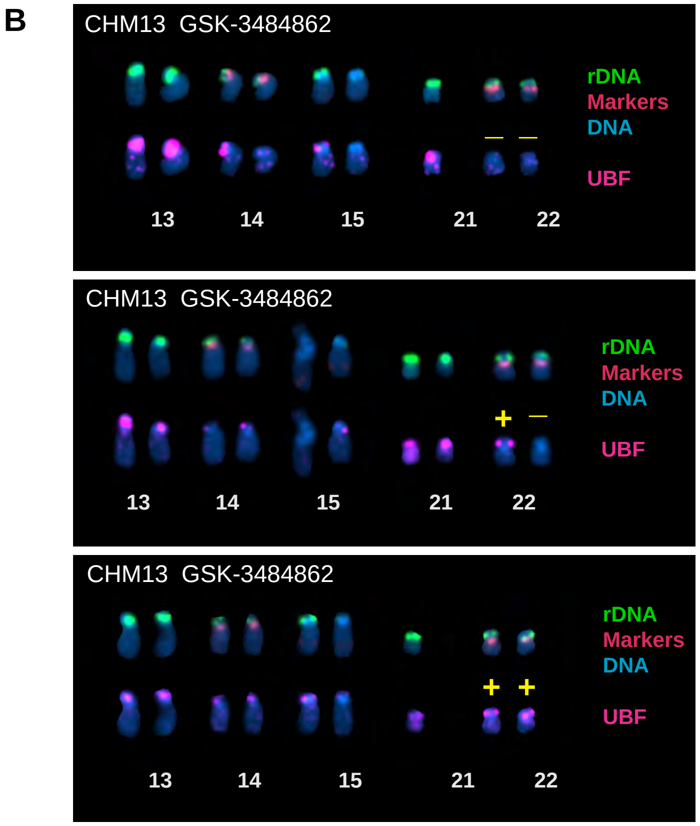


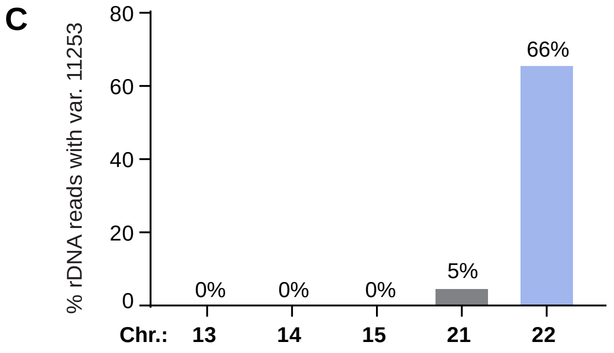


**C.** Fractions of rDNA reads containing variant at the position 11253. Acrocentric chromosomes from CHM13 cells were sorted as in Figure 3A and analyzed by Illumina short-read sequencing using the mpileup tool. 66% of rDNA reads from sorted chromosome 22 contained the variant, compared to 0-5% of reads from other acrocentric chromosomes.

**B.** Extended acrocentric karyograms from DNMT1 inhibitor GSK-3484862 treated CHM13 cells shown in Figure 4C. The top rows of chromosomes show FISH labeling with rDNA probe (green) and chromosome identification markers CenSat 14/22 and PML (red). The bottom rows show corresponding chromosomes with UBF antibody labeling (magenta). DNA was counter-stained with DAPI.

**Supplementary Figure 7. rDNA copy number and activity of HG002 iPS cells and family trio**

A

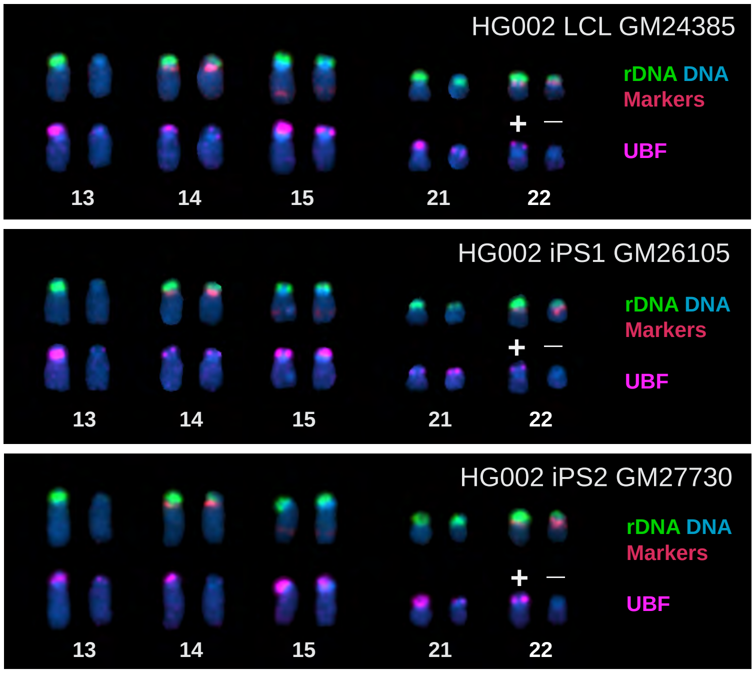


**A.** Extended acrocentric karyograms from HG002 LCL and iPS cell lines.
Top rows of chromosomes show FISH labeling with rDNA probe (green) and chromosome identification markers CenSat 14/22 and PML on chr.15 (red). Bottom rows show corresponding chromosomes with UBF antibody labeling (magenta). DNA was counter-stained with DAPI.

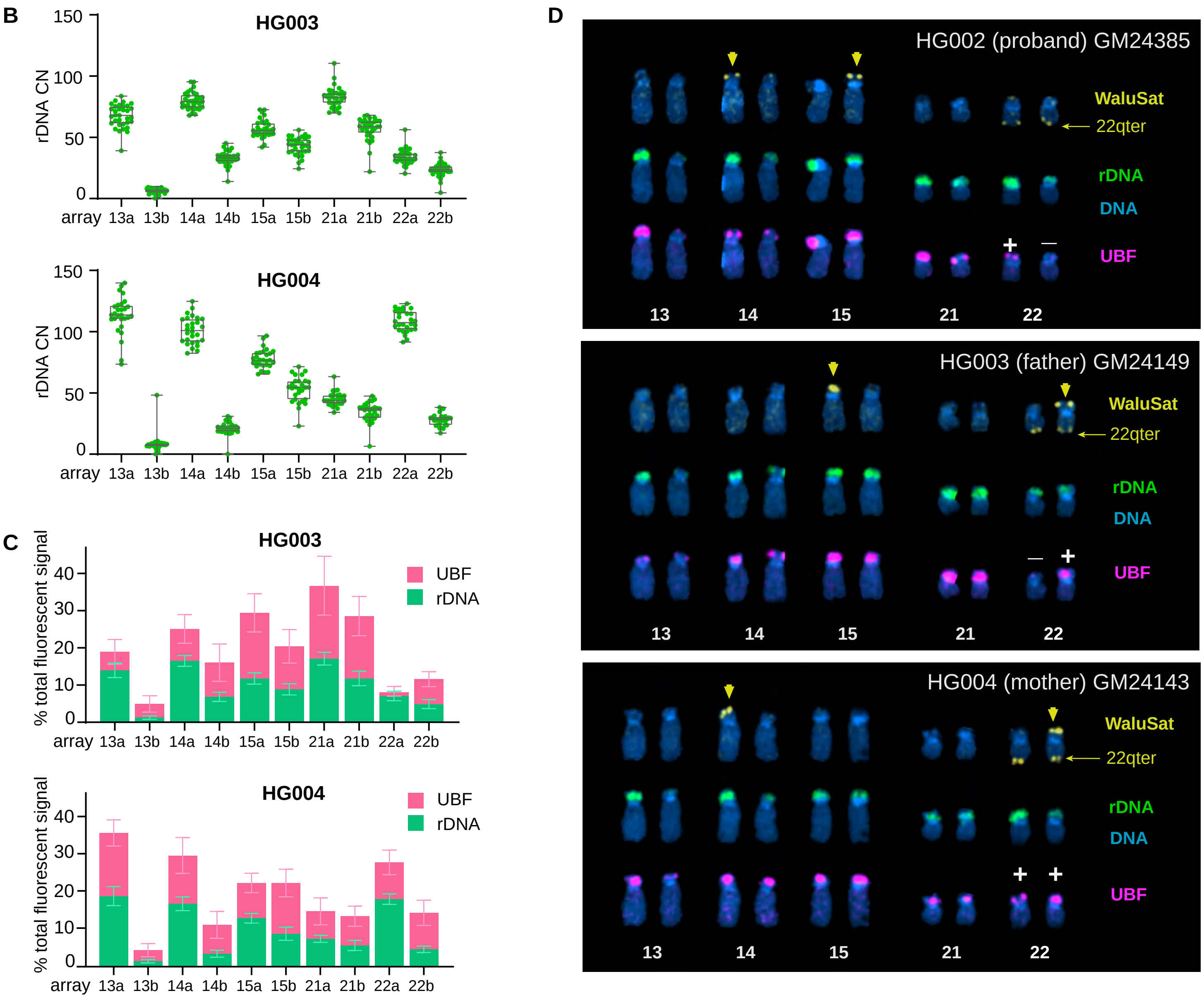

**B.** Quantification of rDNA copy number on acrocentric arrays in HG003 and HG004 LCLs. Chromosome spreads were labeled by FISH with probes for rDNA and chromosome identification markers. The rDNA copy number on each array was calculated from its fraction of the total fluorescent intensity of the rDNA signals on all acrocentric chromosomes and the Illumina sequencing estimate of the total copy number of rDNA repeats in the genome. The boxes represent the interquartile range, with the edges indicating the upper and lower quartiles. The line inside the boxes indicates the median. Whiskers show the range from minimum to maximum values. All individual data points are shown.

**C.** Quantification of rDNA array sizes and activity in HG003 and HG004 LCLs. Chromosome spreads were labeled with rDNA probe, identification markers, and UBF antibody as in panel A. The green and pink sections of the bars represent averages of rDNA and UBF, respectively, expressed as fractions of the total signal in the chromosome spread. Error bars denote standard deviation.

**D.** Extended acrocentric karyograms from HG002, HG003 and HG004 LCL cell lines. Top rows show chromosomes labeled by FISH with WaluSat probe (yellow signal present on some acrocentric p-arms (highlight by arrows) and a chromosome 22 q-terminal region probe (arrow) serving as identification marker for chromosome 22. Middle rows show corresponding chromosomes labeled with rDNA probe (green). Bottom rows show the corresponding chromosomes labeled with UBF antibody labeling (magenta). DNA was counter-stained with DAPI. Note the UBF-negative rDNA array on the WaluSat-negative copy of chromosome 22 in HG002, and the corresponding paternal copy of chromosome 22 in HG003.
